# Aryl Hydrocarbon Receptor Blocks Aging-Induced Senescence in the Liver and Fibroblast Cells

**DOI:** 10.1101/2021.02.25.432074

**Authors:** Ana Nacarino-Palma, Eva M. Rico-Leo, Judith Campisi, Arvind Ramanathan, Jaime M. Merino, Pedro M. Fernández-Salguero

## Abstract

Aging induces progressive organ degeneration and worsening of tissue homeostasis leading to multiple pathologies. Yet, little is known about the mechanisms and molecular intermediates involved. Here, we report that aged aryl hydrocarbon receptor-null mice (*AhR-/-*) had exacerbated senescence and larger numbers of liver progenitor cells. Senescence-associated markers β-galactosidase (SA-β-Gal), p16^Ink4a^ and p21^Cip1^ and genes of the senescence-associated secretory phenotype (SASP) TNF and IL1 were overexpressed in aged *AhR-/-* livers. AhR binding to the promoter of those genes, as shown by chromatin immunoprecipitation, likely had a repressive effect maintaining their physiological levels in *AhR+/+* livers. Furthermore, factors secreted by senescent cells MCP-2, MMP12 and FGF were also produced at higher levels in aged AhR-null livers. Supporting the linkage between senescence and stemness, liver progenitor cells were more abundant in *AhR-/-* mice, which could probably contribute to their increased hepatocarcinoma burden. These roles of AhR are not liver-specific since adult and embryonic AhR-null fibroblasts acquired cellular senescence upon culturing with overexpression of SA-β-Gal, p16^Ink4a^ and p21^Cip1^. Notably, depletion of senescent cells with the senolytic agent navitoclax restored basal expression of senescent markers in *AhR-/-* fibroblasts. Oppositely, senescence promoter palbociclib induced an AhR-null like phenotype in *AhR+/+* fibroblasts. Moreover, doxycycline-induced senescence reduced AhR levels while depletion of p16^Ink4a^-expressing senescent cells restored basal AhR levels in mouse lungs. Thus, AhR is needed to restrict age-induced senescence, and such activity seems to correlate with a more differentiated phenotype and with increased resistance to liver tumorigenesis.

## INTRODUCTION

The aryl hydrocarbon receptor (AhR) is a transcription factor with important roles in toxicology and cell physiology (Fan et al., 2010; Puga et al., 2005; Roman et al., 2018). AhR has a significant role in essential signaling pathways controlling cell cycle control, cell-cell contact, apoptosis, angiogenesis, differentiation and pluripotency (Cho et al., 2004; Ito et al., 2004; Morales-Hernandez et al., 2016; Roman et al., 2009). Deregulation of these pathways, critical for the maintenance of cellular homeostasis, predispose organisms to develop different pathologies including cancer (Fan et al., 2010; Mulero-Navarro and Fernandez-Salguero, 2016; Safe et al., 2013).

Recent findings have uncovered important roles of AhR in cell differentiation and pluripotency, eventually supporting its impact as therapeutic target (Ko and Puga, 2017; Morales-Hernandez et al., 2016; Mulero-Navarro and Fernandez-Salguero, 2016). Previous work from our group demonstrate that AhR adjusts liver and lung regeneration upon injury by controlling proliferation and the expansion of pluripotent stem-like cells (Morales-Hernandez et al., 2017; Moreno-Marin et al., 2017). Accordingly, lack of AhR (*AhR-/-*) induces an undifferentiated and non-polyploid phenotype in the mouse liver that persists from preweaning to adulthood (Moreno-Marín et al., 2018). Moreover, increased numbers of stem cells in *AhR-/-* mouse livers correlates with their higher susceptibility to develop diethylnitrosamine (DEN)-induced hepatocarcinomas (Fan et al., 2010; Moreno-Marin et al., 2017).

In addition to multiple genetic and environmental causes, aging remains a major risk factor in developing cancer (Kruk et al., 2019). During the aging process, biological systems undergo progressive degeneration with the accumulation of molecular changes that compromise and decrease physiological and fertility functions, ultimately leading to loss of homeostasis and elderly pathologies. In addition, impaired cellular and molecular functions during aging may trigger new and aberrant regenerative capacities leading to the appearance of hyperplasic pathologies (Campisi and Robert, 2014).

Most age-related pathologies, degenerative or hyperplastic, are linked to a stress response called senescence. Senescence consists of an irreversible arrest of the cell cycle, changes in chromatin organization and alteration of gene expression patterns (Gorgoulis et al., 2019; Hernandez-Segura et al., 2018). Such phenotype includes secretion of pro-inflammatory cytokines, chemokines, growth factors and proteases, generating a senescence-associated secretory phenotype (SASP) (Campisi and di Fagagna, 2007). The former cellular response to senescence is to halt proliferation of damaged cells, thus evolving as a protective and barrier mechanism against the development of cancer (Campisi and Robert, 2014; Coppe et al., 2008; Rhinn et al., 2019). Moreover, the fact that SASP has multiple paracrine activities suggests that senescence not only prevents cancer but also serves to promote tissue repair and regeneration upon injury (Campisi, 2013; Chiche et al., 2020; Rhinn et al., 2019). In addition, SASP factors can enhance tumorigenesis by promoting proliferation, metastasis and immunosuppression (Wang et al., 2020). A relevant link has been identified between senescence and cellular reprogramming since *in vivo* reprogramming can induce tumor- (Abad et al., 2013; Mosteiro et al., 2016) and aging-associated senescence (Chiche et al., 2020) by mechanisms probably requiring an exacerbated inflammatory status.

An early report revealed that mouse embryo fibroblasts (MEFs) from *AhR-/-* mice reached senescence earlier than *AhR+/+* cells during adipogenic differentiation (Alexander et al., 1998). Later work has shown that AhR reduces smoking-induced inflammation in the lung parenchyma through, among other mechanisms, the control of senescence (Guerrina et al., 2018). In addition, air-born environmental particles can promote cellular senescence through AhR by the production of reactive oxygen species (ROS) causing epigenetic modifications leading to p16^Ink4a^ activation (Ryu et al., 2019).

Here, we report that aged mice lacking AhR expression have a significant increase in hepatic senescence that correlates with an enhanced content of liver progenitor cells and with a larger incidence in hepatocarcinoma burden. Such phenotype involves altered expression of typical senescence and SASP-related markers. AhR deficiency also induced senescence in adult and embryonic fibroblasts, in which its incidence could be modulated by senolytic or senescence promoters in a receptor-dependent manner. Given the recently discovered functional link between senescence and reprogramming, it seems plausible that AhR acts to properly control the balance of both processes during physiological aging. Such relationship may also underline the exacerbated regenerative response developed by AhR-null mice in liver and lung upon injury.

## RESULTS

### AhR depletion increases liver tumor burden with aging

Adult AhR-null mice maintain an undifferentiated and non-polyploid liver phenotype (Moreno-Marin et al., 2018) that could favor the appearance of age-related hepatocarcinogenesis. To address this issue, we examined *AhR+/+* and *AhR-/-* mice aged 15 to 22 months for the presence of liver tumors. Lack of AhR lead to the generation of liver tumors that were evident at 18 months of age and that accounted for a large fraction of the liver at 22 months **(Fig. 1A)**. *AhR-/-* tumors were larger in size, more vascularized and presenting pyknotic nuclei at the histological level that were suggestive of chromatin condensation and replicative blockade **(Fig. 1B)**. The overall tumor incidence reached values close to 45% of the AhR-null mice and remained below 20% of the AhR wild type animals **(Fig. 1C)**. Additionally, spontaneous hepatocarcinogenesis affected close to 80% of *AhR-/-* mice of the C56BL/J genetic background, confirming that AhR depletion favors liver tumor promotion (results not shown). To determine if lower AhR levels are in fact associated to increased liver tumorigenesis, we next analyzed AhR expression during aging. As shown in **Fig. 1D**, AhR mRNA expression drastically decreased in *AhR+/+* livers from 14 months and remained low until at least 22 months of age. Consistently, a similar protein expression profile was found by immunoblotting analysis of liver extracts from both genotypes **(Fig. 1E)**. Treatment with the physiological agonist FICZ significantly increased liver mRNA levels of the canonical target gene *Cyp1a1* in *AhR+/+* young mice. Further, aging markedly reduced the ability of FICZ to induce *Cyp1a1* expression in AhR expressing liver **(Fig. 1F)**, hence supporting that AhR downregulation may have a causal role in liver hepatocarcinogenesis. In agreement with the known role of AhR in controlling pluripotency and stemness (Ko and Puga, 2017; Mulero-Navarro and Fernandez-Salguero, 2016), enhanced hepatocarcinogenesis in AhR-null liver correlated with their higher numbers of liver progenitor/stem-like cells in both young and aged animals (**Fig. 1G)**. Interestingly, AhR deficiency also produced an increase in bone marrow progenitor cells regardless of the age of mice **(Fig. 1H)**, thus confirming the role of this receptor in maintaining the population of undifferentiated cells in different organs. Enhanced tumor burden in *AhR-/-* mouse liver correlated with increased expression of the glucose transporter Glut4 not only in old by also in young mice **(Supplementary Fig. 1A)**. Accordingly, glucose uptake, as determined by the level of hexokinase activity, was significantly higher in old AhR-null than in *AhR+/+* mice **(Supplementary Fig. 1B)**. Thus, aging in *AhR-/-* mice induces a preferred glycolytic metabolism that could impact their greater susceptibility to develop liver cancer and that correlates with the Warburg effect.

**Figure 1.**
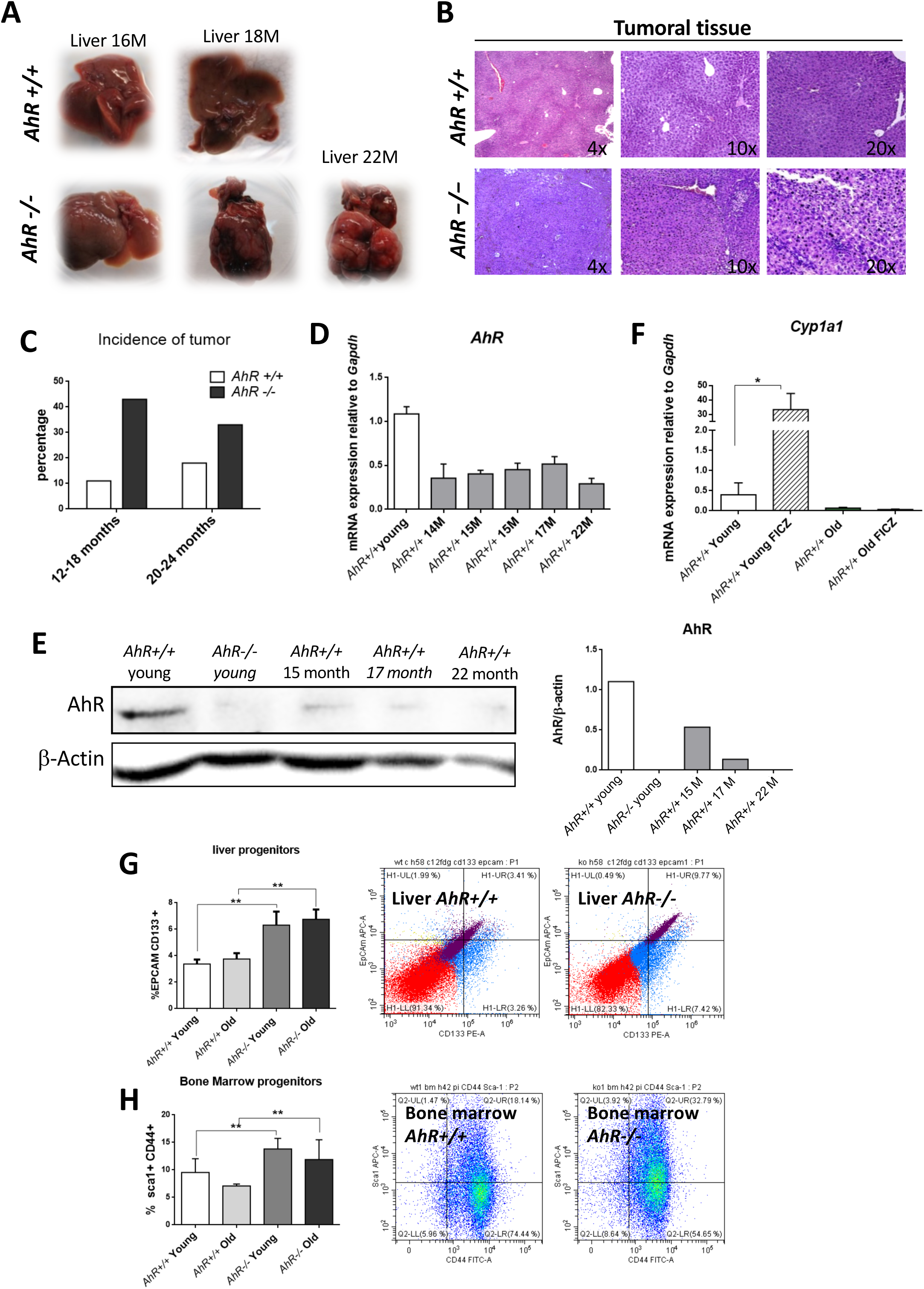
AhR depletion increases liver tumorigenesis with aging. **(A)** Representative tumors developed by *AhR+/+* and *AhR-/-* at the indicated ages. **(B)** Haematoxylin and Eosin staining of liver tumor sections from *AhR+/+* and *AhR-/-* mice at 22 months of age. Note the abundance of pycnotic nuclei in *AhR-/-* tumors. **(C)** Quantification of the number of liver tumors in mice of both genotypes at two different age intervals. **(D)** AhR mRNA expression in *AhR+/+* livers at the indicated ages using RT-qPCR and the oligonucleotides indicated in Supplementary Table 1. **(E)** AhR protein levels were analyzed in liver extracts at the indicated ages by immunoblotting. β-Actin was used to normalize protein levels. **(F)** *AhR+/+* mice were injected i.p. with 4 mg/kg FICZ and the mRNA levels of the AhR canonical target gene *Cyp1a1* were determined by RT-qPCR using the oligonucleotides indicated in Supplementary Table 1. **(G)** Liver progenitor cells were analyzed by FACS using specific antibodies against CD133-PE and EPCAM-APC. Distribution of cell subpopulations and gating from representative experiments are shown. **(H)** Bone marrow progenitor cells were analyzed by FACS using the specific markers CD44-FITC and Sca1-APC. Distribution of cell subpopulations and gating from representative experiments are shown. *Gapdh* was used to normalize target gene expression (ΔCt) and 2^−ΔΔCt^ to calculate changes in mRNA levels with respect to wild type or untreated conditions. Data are shown as mean + SD (\**P <0.05*; \*\**P<0.01*).

### AhR deficient livers have increased senescence with aging

An undifferentiated status favors hepatocarcinogenesis (Kuo et al., 2016; Yin et al., 2015) whereas senescence seems to have an important role in the fate of aged undifferentiated tumors (Chiche et al., 2020; Mosteiro et al., 2018). We then investigated whether AhR deficiency in aged liver could induce a senescent status eventually linked to the generation of hepatic tumors. Senescence-associated β-galactosidase activity (SA-β-Gal) was markedly increased in liver sections from aged *AhR-/-* mice as compared to their *AhR+/+* counterparts **(Fig. 2A)**. Quantification of senescence in C12FDG-labelled hepatic cells by FACS confirmed that AhR deficiency significantly expanded the number of senescent cells upon aging **(Fig. 2B)**. Next, we used liver mRNA to analyze by RT-qPCR the expression of genes considered drivers of senescence. The results showed that, with respect to *AhR+/+* livers, *p16^Ink4a^* mRNA levels markedly increased in *AhR-/-* livers at 15 months to decrease to lower levels at 22 months of age **(Fig. 2C)**. Similarly, *p21^Cip1^* transiently increased in *AhR-/-* livers at 15 months reaching values significantly higher to those of *AhR+/+* livers; differences in *p21^Cip1^* expression between both genotypes could be seen even in young mice **(Fig. 2D)**. The expression of markers involved in the senescence-associated secretory phenotype (SASP) IL1 and TNFα was also up-regulated with aging in absence of AhR **(Fig. 2E,F)**, although TNFα remained at higher levels even at the oldest age analyzed (22 months). We did not detect significant transcriptional changes between both genotypes with respect to liver p53 or IL6 (data not shown). Thus, AhR limits senescence and carcinogenesis in the liver and such activities could be related to its potential to maintain differentiation in this organ.

**Figure 2.**
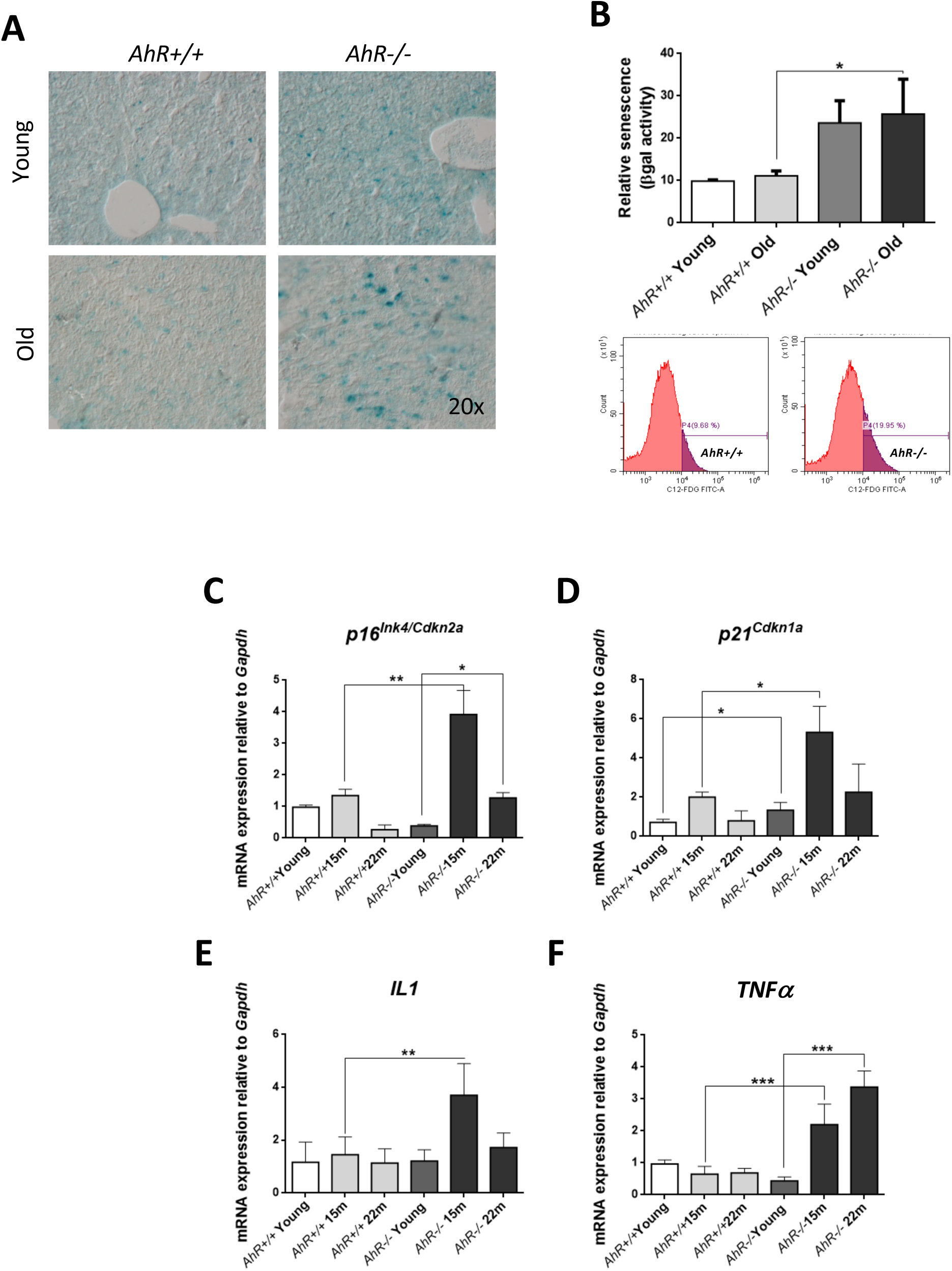
Cell senescence increases with age in AhR-deficient liver. **(A)** SA-β-Gal activity was analyzed in *AhR+/+* and *AhR-/-* liver sections by staining with the chromogenic substrate X-Gal. **(B)** SA-β-Gal activity was analyzed by FACS in isolated liver cells (gentleMACS) using the fluorescent substrate C12FDG. **(C-F)** mRNA expression levels of senescence driver genes *p16^Ink4a^ (***C)** and *p21^Cip1^* **(D)** and SASP-related genes IL1 **(E)** and TNFα **(F)** were analyzed by RT-qPCR in *AhR +/+* and *AhR-/-* livers at the indicated ages. Oligonucleotides used are indicated in Supplementary Table 1. *Gapdh* was used to normalize target gene expression (ΔCt) and 2^−ΔΔCt^ to calculate changes in mRNA levels with respect to wild type or untreated conditions. Data are shown as mean + SD (\**P <0.05*; \*\**P<0.01;* \*\*\**P<0.001*).

To further correlate undifferentiation and senescence in the liver, we analyzed markers of both processes in non-tumoral tissue from old mice and in hepatocarcinoma samples from *AhR+/+* and *AhR-/-* mice by immunofluorescence. Regarding senescence, p16^Ink4a^ and p21^Cip1^were present at higher levels in the liver of aged AhR-null mice as compared to wild mice. Pluripotency/stemness and undifferentiation inducers NANOG and OCT4 were also overexpressed in *AhR-/-* livers of old mice **(Fig. 3)**. Expression of p21^Cip1^ and OCT4 appeared to be prevalent with respect to p16^Ink4^ and NANOG, respectively. Well-developed hepatocarcinomas from *AhR-/-* mice **(Fig. 1)** showed even higher levels of each of those proteins, suggesting that tumorigenesis accentuate the basal undifferentiated and senescent status of aged *AhR-/-* mice. The mesenchymal marker α-smooth muscle actin (α-SMA) had increased levels in hepatocarcinomas and old *AhR-/-* liver as compared to livers of *AhR+/+* mice **(Fig. 3)**, in agreement with the known accumulation of extracellular matrix in old AhR-null liver (Fernandez-Salguero et al., 1995; Schmidt et al., 1996) that may eventually contribute to hepatocarcinogenesis.

**Figure 3.**
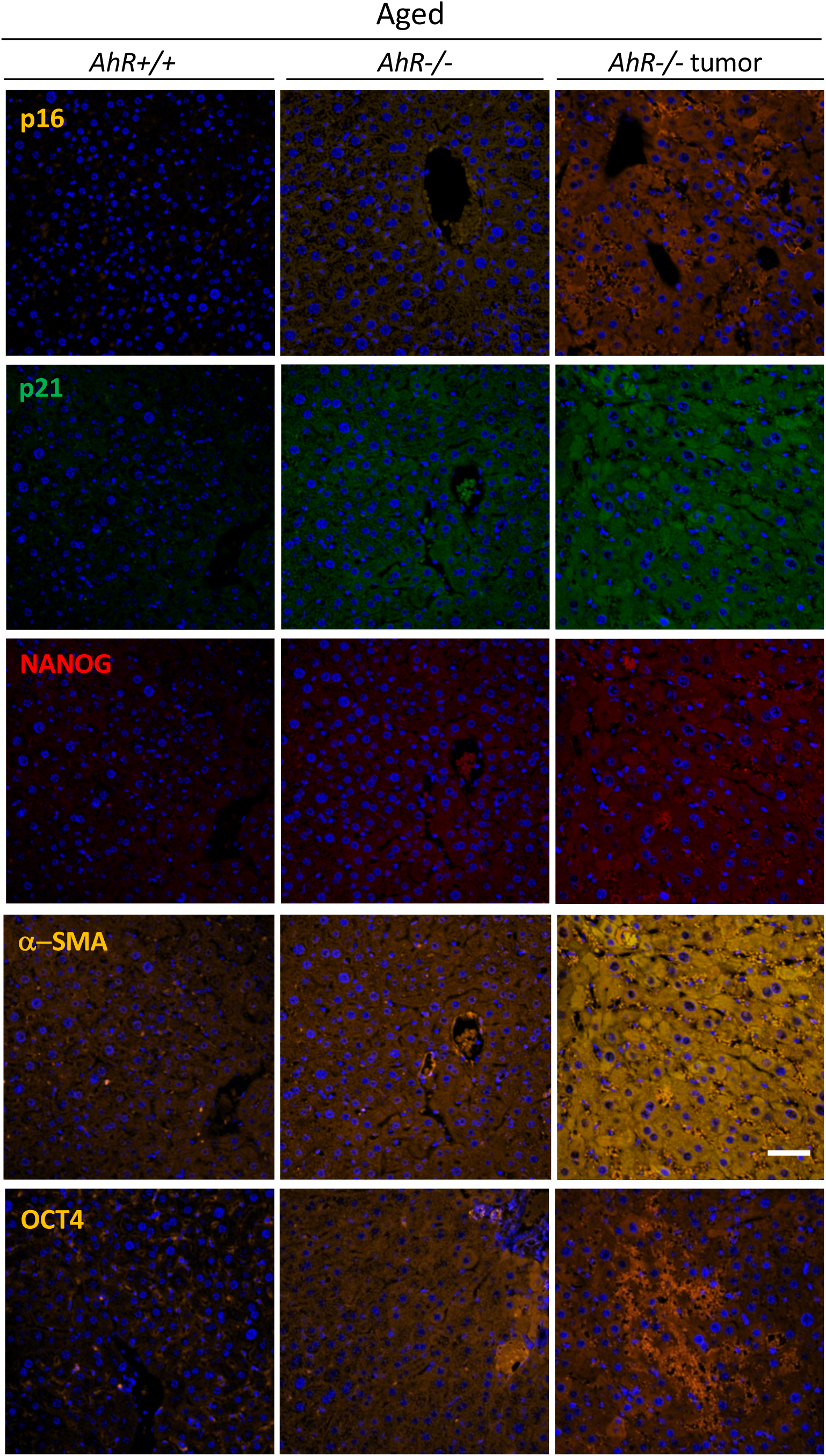
*AhR-/-* livers overexpress senescence and undifferentiation/stemness markers. Protein levels for senescence markers p16^Ink4^ and p21^Cip1^ and for pluripotency/stemness inducers NANOG and OCT4 were analyzed by immunofluorescence in liver sections of aged *AhR+/+* and *AhR-/-* mice and in hepatocarcinomas from *AhR-/-* mice using specific antibodies. The expression of α-SMA was also analyzed as indicator of vasculogenesis. Conjugated secondary antibodies labelled with Alexa 488, Alexa 550 and Alexa 633 were used for detection. DAPI staining was used to label cell nuclei. An Olympus FV1000 confocal microscope and the FV10 software (Olympus) were used for the analysis. Scale bar corresponds to 50 μm.

Since senescence markers were up-regulated at the mRNA level in absence of AhR **(Fig. 2)**, we next decided to analyze if AhR transcriptionally represses those genes by direct binding to their promoters. Sequence analysis revealed the presence of the AhR-canonical binding site XRE (Xenobiotic response element, 5’GCGTG 3’) in the upstream promoter region of *p16^Ink4a^, p21^Cip1^* and *TNFα.* Chromatin immunoprecipitation (ChIP) for AhR in liver extracts from young mice (2 months) and mice aged 15 and 22 months was performed **(Fig. 4)**. AhR was bound to the *p16^Ink4a^* gene promoter at 15 months to be released in older animals at 22 months **(Fig. 4A)**. AhR binding to the *p21^Cip1^* and *TNFα* gene promoters was maximal at 22 months of age and nearly basal in younger mice **(Fig. 4B,C)**. Thus, overexpression of these genes in old *AhR-/-* livers likely results from a lack of transcriptional repression in absence of receptor.

**Figure 4.**
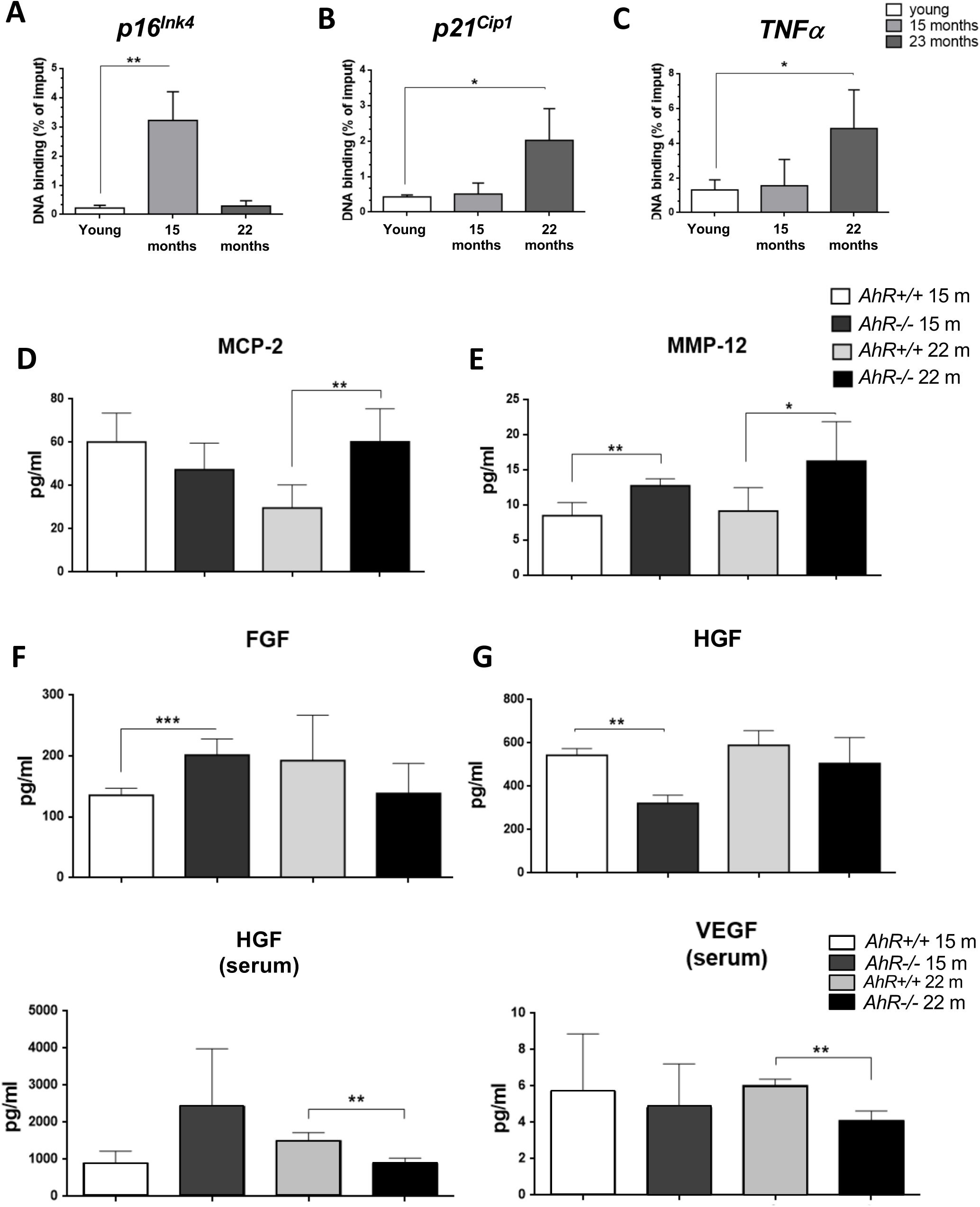
AhR modulates the senescence-associated secretory phenotype with aging. **(A).** Chromatin immunoprecipitation (ChIP) for AhR binding to XRE binding sites located in the promoter of *p16^Ink4a^* **(A)**, *p21^Cip1^* **(B)** and *TNFα* **(C)**. qPCR was used to quantify changes in DNA binding and the results were normalized to the corresponding inputs. Amounts of MCP-2 **(D)**, MMP12 **(E)**, FGF **(F)** and HGF **(G)** were analyzed in liver homogenates from *AhR+/+* and *AhR-/-* mice at the indicated ages. Levels of HGF **(H)** and VEGF **(I)** VEGF were also determined in serum from mice of the same genotypes and ages. Bio-Plex Multiplex immunoassays kit were used. Oligonucleotides for qPCR are indicated in Supplementary Table 1. *Gapdh* was used to normalize target gene expression (ΔCt) and 2^−ΔΔCt^ to calculate changes in mRNA levels with respect to wild type or untreated conditions. Data are shown as mean + SD (\**P<0.05*; \*\**P<0.01;* \*\*\**P<0.001*).

### Factors of the senescence-associated secretory phenotype are regulated in an AhR-dependent manner during liver aging

Soluble factors secreted by senescent cells could contribute to the aging phenotype of *AhR-/-* mouse liver. To address that, we used immunoassays to analyze the levels of major SASP factors secreted within the liver or circulating in the blood of *AhR+/+* and *AhR-/-* aged mice. In the liver, secretion of MCP2/CCL8, a cytokine responsible for chemoattraction of monocytes and macrophages to damaged or tumor areas, was significantly higher in *AhR-/-* mice at 22 months as compared to age-matched *AhR+/+* mice **(Fig. 4D).** Extracellular matrix metalloproteinase MMP-12, a stimulator of matrix degradation and liver tumor progression, was significantly overexpressed not only in the oldest (e.g., 22 months) but also in mice at the beginning of the aging process (e.g., 15 months) **(Fig. 4E)**. Moreover, fibroblast growth factor (FGF), whose expression seems to inhibit senescence (Coutu and Galipeau, 2011), had a trend to decrease in *AhR-/-* liver with aging, despite having higher levels than wild type mice at 15 months **(Fig. 4F)**. Liver hepatocyte growth factor (HGF), whose overactivation has been related to senescence inhibition (Zhu et al., 2014), was also reduced at 15 months and not significantly increased in older 22 months AhR-null mice **(Fig. 4G)**. In serum, HGF levels were significantly lower in oldest 22 months *AhR-/-* mice **(Fig. 4H),** as it was the amount of circulating vascular endothelial growth factor (VEGF) **(Fig. 4I)**, in agreement with its implication in aging-related endothelial senescence (Regina et al., 2016). Thus, the SASP appears to be exacerbated in *AhR-/-* liver during aging.

### Cell senescence markers are enhanced in adult primary fibroblasts lacking AhR

To investigate if the role of AhR in senescence is extensive to additional cell types, we decided to examine adult primary fibroblasts as they are known targets of senescence (Alvarez et al., 2017; Zorin et al., 2017) and can be readily isolated from adult *AhR+/+* and *AhR-/-* tissues. Adult tail tip fibroblasts (TTFs) from young (4-6 weeks) and old mice (15-22 months) of both genotypes were isolated, cultured and analyzed for their senescence level using the SA-β-Gal fluorescent substrate C12FDG **(Fig. 5).** Fluorescent confocal microscopy revealed that both young and old *AhR-/-* TTFs had larger numbers of senescent cells than *AhR+/+* TTFs grown under the same culturing conditions **(Fig. 5A)**. Flow cytometry similarly showed that lack of AhR significantly increased the population of senescent cells in *AhR-/-* TTFs at both ages, as revealed by C12FDG fluorescence **(Fig. 5B)**. Cells positive for p16^Ink4a^ were equally abundant in young TTFs of both genotypes but markedly increased in old *AhR-/-* cells **(Fig. 5C)**. Regarding p21^Cip1^ a similar response was found in *AhR-/-* TTFs with aging, being its expression nuclear in a large fraction of cells **(Fig. 5D)**. On the other hand, old *AhR-/-* TTFs had a larger number of Cyclin E positive cells than old *AhR+/+* TTFs **(Fig. 5E)**, together with a higher percentage of cells at the G0/G1transition of the cell cycle that was also present in young AhR-null cells **(Fig. 5F)**. These results suggest that a concurrent increase in Cyclin E and tumor suppressors may cause cell cycle blockade, reduced proliferation of adult *AhR-/-* fibroblasts (Mulero-Navarro et al., 2005; Zaher et al., 1998) and eventually contribute to higher rates of senescence.

**Figure 5.**
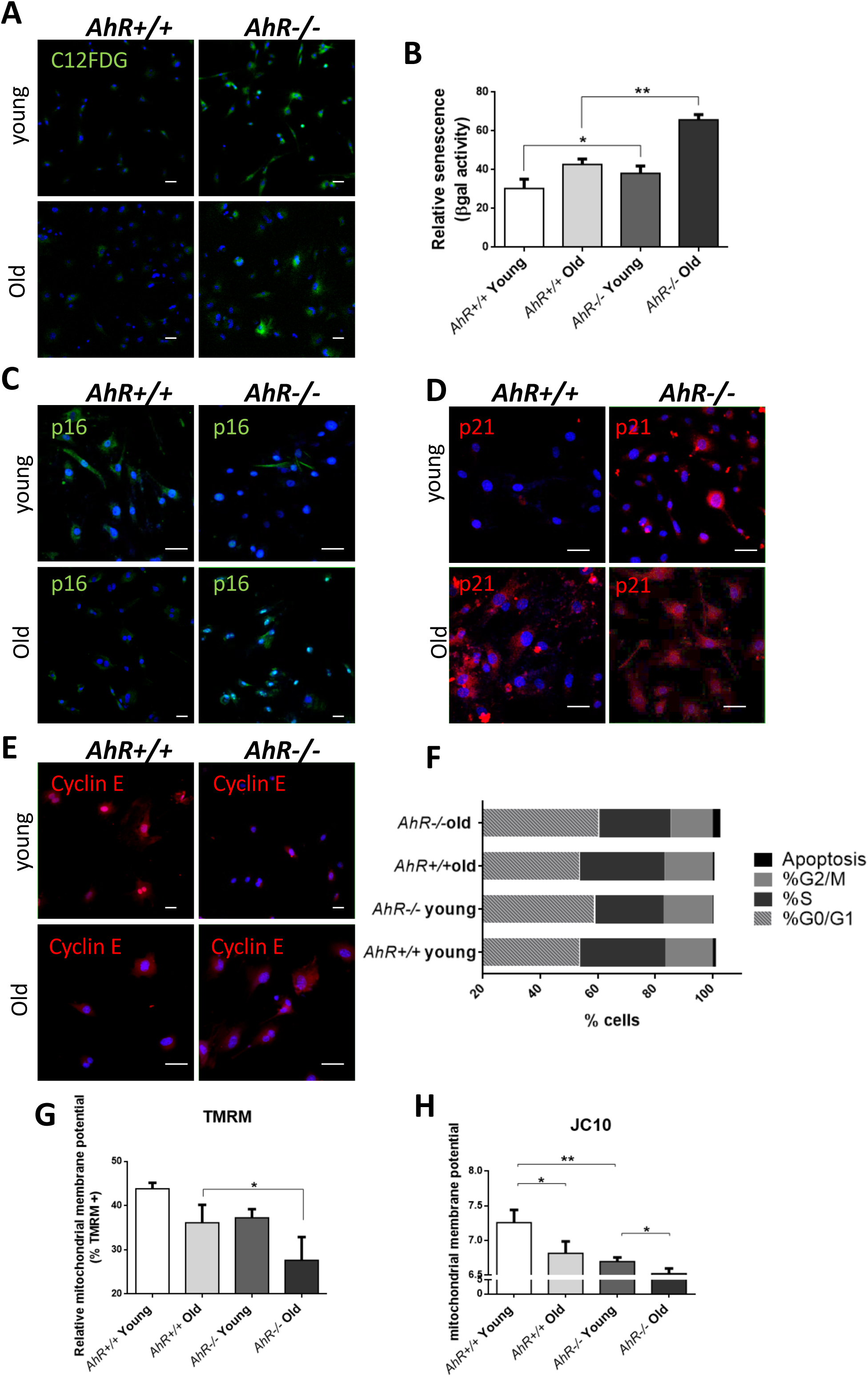
Senescence increases with aging in adult AhR-null fibroblasts. **(A)** *AhR+/+* and *AhR-/-* TTFs were stained with the SA-β-Gal fluorescent substrate C12FDG to determine senescence levels by confocal microscopy. **(B)** SA-β-Gal activity was also measured by flow cytometry analyzing the percentage of C12FDG positive cells. **(C-E)** p16^Ink4a^ **(C),** p21^Cip1^ **(D)** and Cyclin E **(E)** were analyzed by florescent confocal microscopy using specific antibodies in young and aged fibroblast cells. Conjugated secondary antibodies used were Alexa 488 and Alexa 633. DAPI staining was used to label cell nuclei. An Olympus FV1000 confocal microscope with FV10 software (Olympus) were used. **(F)** Percentage of TTFs in each cell cycle phase. Cell cycle analysis was performed by FACS using propidium iodide staining. **(G)** Mitochondrial membrane potential (MMP) was quantified by the percentage of TMRM positive cells analyzed by Cytoflex S cytometer (Beckman Coulter). **(H)** MMP was also calculated using the JC10 kit for mitochondrial membrane polarization (Sigma-Aldrich). The red/green fluorescence intensity ratio was used to determine MMP activity. Data are shown as mean + SD (\**P <0.05*; ** *P <0.01*).

Mitochondrial dysfunction is an additional parameter indicative of cellular senescence (Moro, 2019). We thus measured the mitochondrial membrane potential (MMP) of young and old *AhR+/+* and *AhR-/-* TTFs. Flow cytometry analyses using tetramethyl rhodamine (TMRM), a fluorescent cationic dye captured by active mitochondria, showed that old AhR-null TTFs had reduced mitochondrial activity than *AhR+/+* cells **(Fig. 5G).** We latter used the JC10 probe, which forms aggregates depending on membrane polarization, to further support that old *AhR-/-* mice had lower mitochondrial metabolism **(Fig. 5H)**. Therefore, aged primary adult fibroblasts have increased senescence rates, reduced cell cycling and mitochondrial dysfunction suggestive of glycolytic metabolism.

### Embryonic primary fibroblasts lacking AhR expression have increased senescence rates

AhR-null mouse embryonic fibroblasts (MEF), on *in vitro* passaging, acquire a senescent phenotype earlier than wild type cells (Alexander et al., 1998) and have reduced proliferation potential (Santiago-Josefat et al., 2004). Flow cytometry and cytological staining for SA-β-Gal indicated that, upon *in vitro* culturing, AhR-null MEFs had higher levels of senescence than *AhR+/+* MEFs, which became significant from passage 3 (P3) **(Fig. 6A-C)**. AhR expression gradually increased with the number of passages from P2 to P5 **(Fig. 6D)**, a result suggesting that, in undifferentiated *AhR+/+* embryonic cells, AhR up-regulation may be a response to restrict senescence and that full receptor depletion may exacerbate such process. Consistently, senescence drivers *p16^Ink4a^* and *p21^Cip1^* were up-regulated in *AhR-/-* MEFs with the number of passages following a pattern closely to that of accumulation of senescent cells **(Fig. 6E,F)**. Levels of p53 did not significantly change during in vitro passaging in either genotype **(Fig. 6G)**.

**Figure 6.**
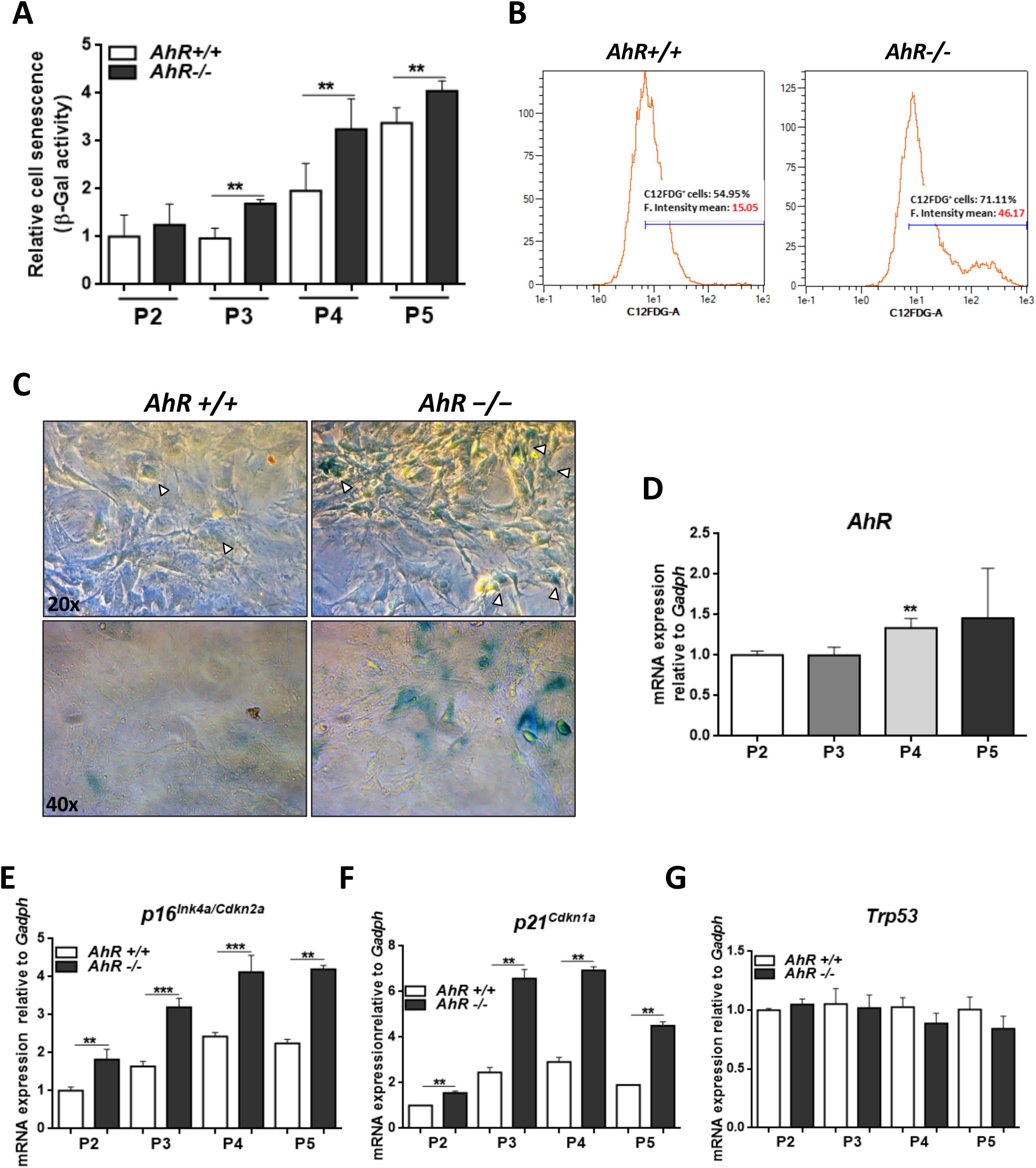
Lack of AhR enhances *in vitro* cellular senescence in mouse embryonic fibroblasts. **(A)** Senescence profile of *AhR+/+* and *AhR-/-* MEFs at the indicated passages as determined by the level of SA-β-gal activity analyzed by FACS using the β-galactosidase fluorescent substrate C12FDG. Results are normalized to wildtype MEFs at passage 2. **(B)** Representative flow cytometric profiles of senescent cells stained with C12FDG in *AhR+/+* and *AhR-/-* MEFs at passage 5. **(C)** SA-β-Gal activity in *AhR+/+* and *AhR-/-* MEFs at passage 5 as determined by cellular staining using X-Gal as substrate. **(D)** *AhR* mRNA expression was determined by RT-qPCR at the indicated cell culture passages. **(E-G)** mRNA expression of senescence driver genes *p16^Ink4a^* **(E)***, p21^Cip1^* **(F)** and *Trp53* **(G)** were determined in *AhR+/+* and *AhR-/-* MEFs by RT-qPCR using the oligonucleotides indicated in Supplementary Table 1. *Gapdh* was used to normalize target gene expression (ΔCt) and 2^−ΔΔCt^ to calculate changes in mRNA levels with respect to wild type or untreated conditions. Data are shown as mean + SD (\*\**P<0.01;* \*\*\**P<0.001*).

To further characterize the contribution of AhR to cellular senescence, we choose to treat MEF cells with specific inhibitors and activators of the process. Bcl-2 family inhibitor Navitoclax is a senolytic agent able to block cell senescence (Zhu et al., 2016). Navitoclax significantly reduced senescence rates in *AhR+/*+ and *AhR-/-* MEFs with respect to untreated control cultures **(Fig. 7A-C)**. In agreement with the up-regulation of AhR in wild type senescent cells **(Fig. 6D)**, Navitoclax induced a decrease in receptor levels **(Fig. 7D)** that again suggest a coordinated regulation between its expression and the magnitude of senescence. As expected, this senolytic compound significantly reduced the expression of senescence markers *p16^Ink4a^* and *p21^Cip1^* in *AhR-/-* MEFs as compared with vehicle-treated cultures **(Fig. 7E,F)**. However, Navitoclax appeared to increase p53 levels in AhR deficient MEF cells **(Fig. 7G)**.

**Figure 7.**
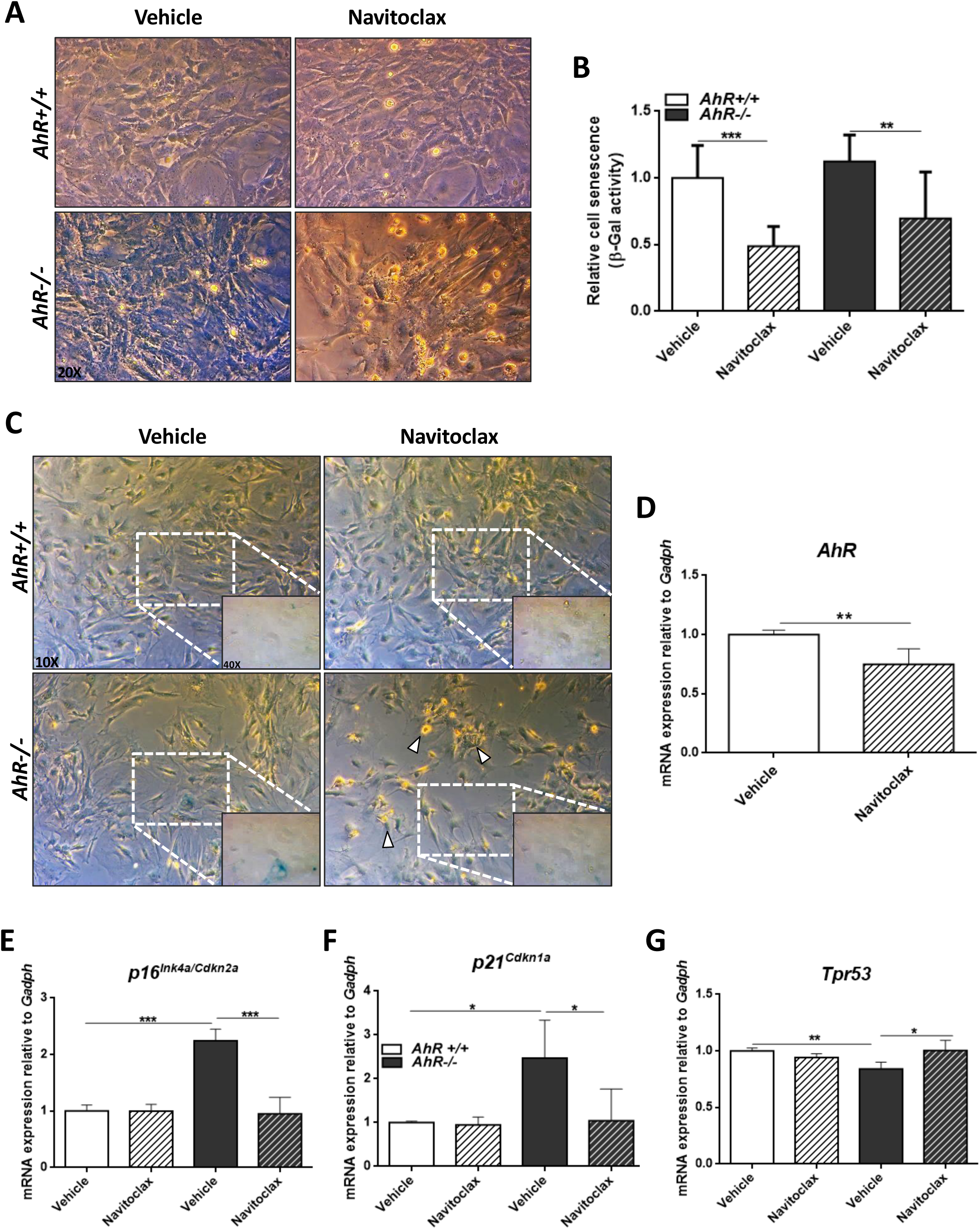
Senolytic agent Navitoclax restores wild-type mRNA levels of senescence driver genes in *AhR-/-* MEFs. Embryonic fibroblast cultures at P4 or P5 were treated with vehicle or 10 μM Navitoclax for 48 h. (**A)** Bright-field microscopy of *AhR+/+* and *AhR-/-* MEFs untreated or treated with Navitoclax. (**B)** Cell senescence measured as percentage of SA-β-Gal activity in MEF cells of both genotypes. SA-β-Gal activity was analyzed by FACS using the β-galactosidase fluorescent substrate C12FDG. Results are normalized to vehicle-treated wild-type MEFs. (**C)** X-Gal staining in untreated and Navitoclax-treated *AhR+/+* and *AhR-/-* MEFs. (**D)** *AhR* mRNA expression was determined by RT-qPCR in both experimental conditions using the oligonucleotides indicated in Supplementary Table 1. **(E)** mRNA expression of senescence driver genes was determined in *AhR+/+* and *AhR-/-* MEFs by RT-qPCR using the oligonucleotides indicated in Supplementary Table 1. *Gapdh* was used to normalize target gene expression (ΔCt) and 2^−ΔΔCt^ to calculate changes in mRNA levels with respect to wild type or untreated conditions. Data are shown as mean + SD (\**P<0.05;* \*\**P<0.01;* \*\*\**P<0.001*).

CDK4/6 inhibitor Palbociclib/PD0332991 has been identified as a cell cycle inhibitor and senescence inducer (Yoshida et al., 2016). Palbociclib efficiently increased senescence rates in both *AhR+/+* and *AhR-/-* MEFs although its effects were more pronounced in the former due to their lower basal incidence of senescent cells **(Fig. 8A-C)**. Treatment of wild type MEFs at passage 2 with Palbociclib for 8 days did not alter AhR expression with respect to solvent-treated control cultures **(Fig. 8D)**, perhaps because under those non-proliferating and confluent conditions *AhR+/+* MEFs normalize AhR expression regardless of the levels of senescence. In agreement with its senescence induction activity, Palbociclib significantly increased *p16^Ink4a^* and *p21^Cip1^* expression in MEFs of both genotypes, despite a less accused effect in *AhR-/-* MEFs likely due to their higher basal level of both proteins **(Fig. 8E,F)**. The expression of p53 was not altered by Palbociclib in either genotype **(Fig. 8G)**.

**Figure 8.**
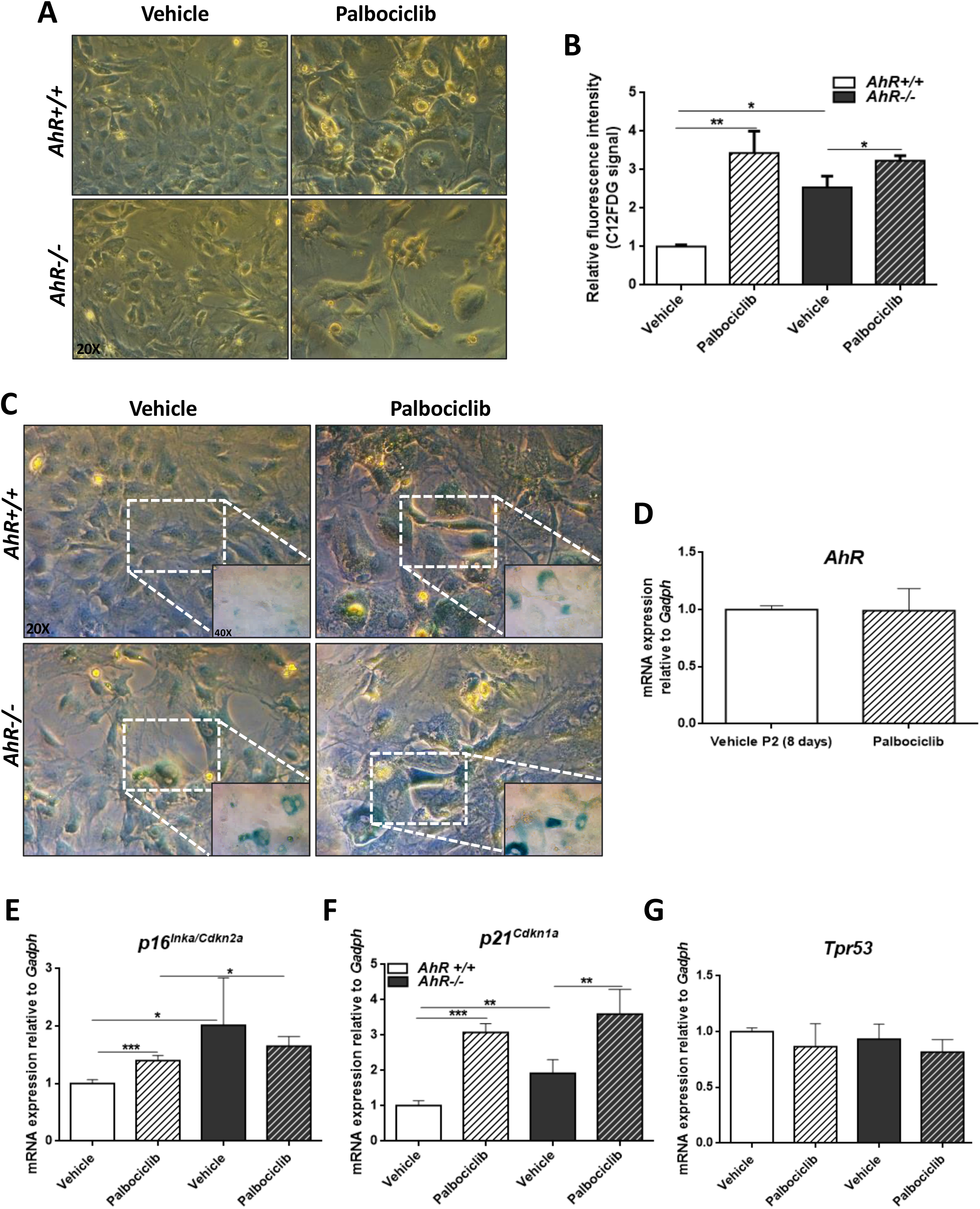
CDK4/6 inhibitor Palbociclib induces senescence in MEF cells and mimicks AhR deficiency. Embryonic fibroblasts were grown in culture for 48 h and then plated at a density of 4 x10^5^ cells per 10-cm plate. Two days later, cells were treated with 4 μM of CDK4/6 inhibitor Palbociclib/PD-0332991 for 8 days. (**A)** Bright-field microscopy of *AhR+/+* and *AhR-/-* MEFs untreated or treated with Palbociclib. (**B)** Cell senescence measured as SA-β-Gal activity in *AhR+/+* and *AhR-/-* MEFs. SA-β-Gal activity was analyzed by FACS using the β-galactosidase fluorescent substrate C12FDG. Results are normalized to vehicle-treated wild-type MEFs. (**C)** X-gal staining in untreated and Palbociclib treated *AhR+/+* and *AhR-/-* MEFs. (**D)** *AhR* mRNA expression was determined by RT-qPCR in *AhR+/+* MEFs under both experimental conditions using the oligonucleotides indicated in Supplementary Table 1. Determinations were done after 8 days of treatment with Palbociclib or with vehicle control. (**E)** mRNA expression of senescence driver genes was determined in *AhR+/+* and *AhR-/-* MEFs by RT-qPCR using the oligonucleotides indicated in Supplementary Table 1. *Gapdh* was used to normalize target gene expression (ΔCt) and 2^−ΔΔCt^ to calculate changes in mRNA levels with respect to wild type or untreated conditions. Data are shown as mean + SD (\**P<0.05;* \*\**P<0.01;* \*\*\**P<0.001*).

The correlation between AhR expression and induction of senescence in adult tissues was also analyzed *in vivo* using the *p16^Ink4a^-3MR* transgenic mice in which senescent cells can be induced by treatment with doxycycline and eliminated through apoptosis by the metabolism of ganciclovir driven by the *p16^Ink4a^* promoter (Demaria et al., 2014) **(Fig. 9A)**. Senescence induction by doxycycline reduced AhR expression in *p16^Ink4a^-3MR* mice lungs (taken as target tissue) whereas co-treatment with ganciclovir restored receptor levels **(Fig. 9B)**. Accordingly, levels of *p16^Ink4a^* and *p21^Cip1^* were upregulated by doxycycline to be reduced after the elimination of senescent cells by ganciclovir **(9Fig. 9C,D)**. In addition, the expression of *p53* and the SASP factor *Mmp3* was also modulated following induction or elimination of senescent cells **(Fig. 9E,F).**

**Figure 9.**
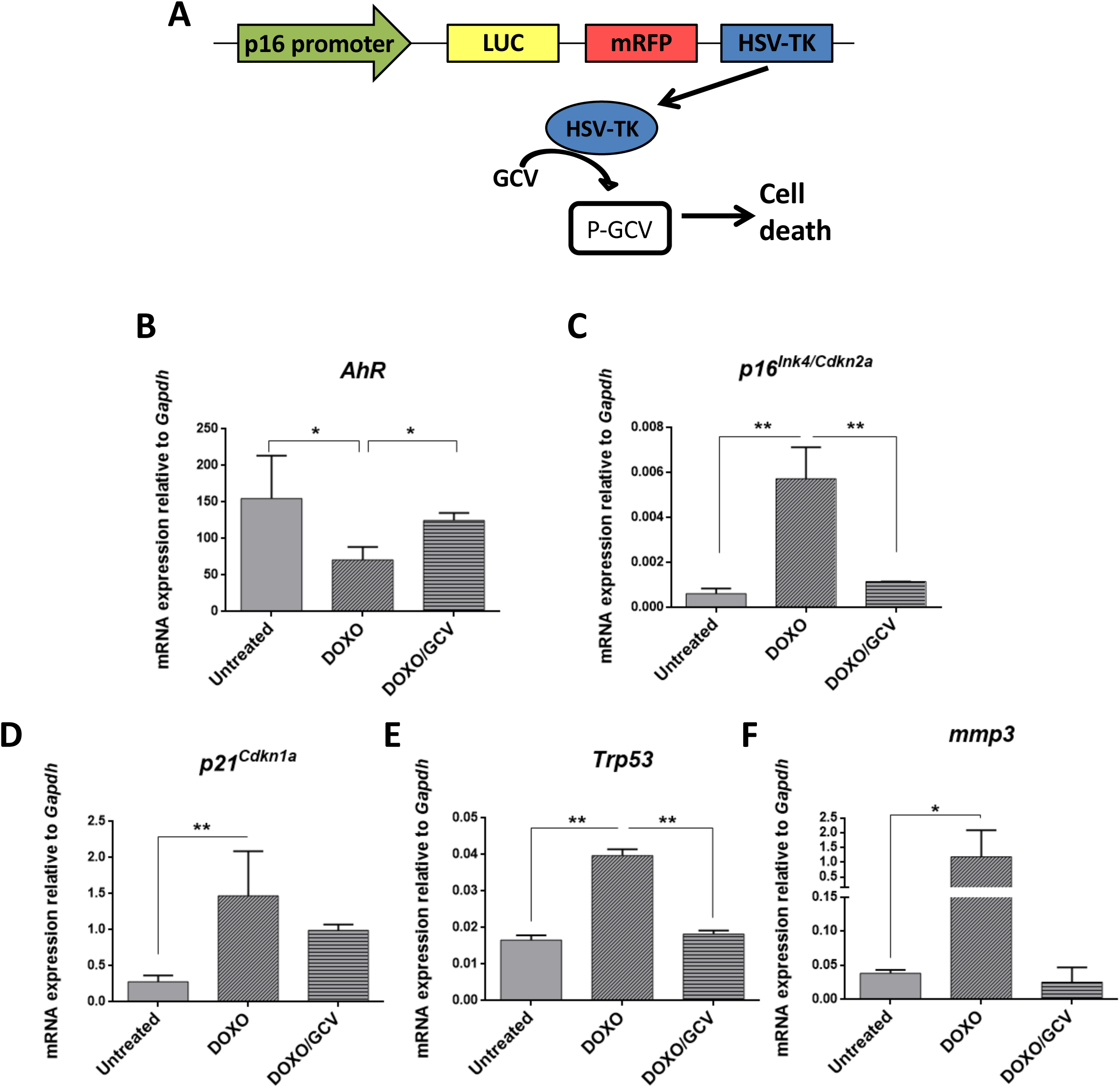
AhR expression varies with the eradication of senescence *in vivo* in *p16^Ink4a^-3MR* transgenic mice. **(A)** Schematic representation of the construct used to generate the *p16^Ink4a^-3MR* transgenic mice to deplete senescent cells *in vivo* by ganciclovir treatment. *p16^Ink4a^-3MR* mice at 4-7 months of age were treated with doxycycline (DOXO), doxycycline + ganciclovir (DOXO+GCV) or vehicle PBS (vehicle). mRNA expression of *AhR* **(B)***, p16^Ink4a^* **(C)**, *p21^Cip1^* **(D),** *p53* **(E)** and Mmp3 **(F)** was determined by RT-qPCR in lung tissue using the oligonucleotides indicated in Supplementary Table 1. *Gapdh* was used to normalize target gene expression (ΔCt) and 2^−ΔΔCt^ to calculate changes in mRNA levels with respect to wild type or untreated conditions. Data are shown as mean + SD (\**P<0.05;* \*\**P<0.01*).

## DISCUSSION

Aging is one major life-threatening condition that increases the incidence of elderly-related diseases including cardiovascular, oncogenic and neurodegenerative pathologies negatively affecting life expectancy. An increasing number of studies are reporting different functions of AhR in aging (Bravo-Ferrer et al., 2019; Brinkmann et al., 2019; Eckers et al., 2016; Kaiser et al., 2020), having a positive or negative role depending on specie, cell phenotype and receptor levels (Mulero-Navarro and Fernandez-Salguero, 2016; Puga et al., 2009; Roman et al., 2018; Safe, 2001). On the other hand, AhR has emerged as a regulatory factor in pluripotency and stemness in different cell types including undifferentiated human teratoma cells (Ko and Puga, 2017; Ko et al., 2013; Morales-Hernandez et al., 2016). Accordingly, this receptor is required to block hepatocarcinogenesis induced by the toxic compound diethyl nitrosamine (Fan et al., 2010; Moreno-Marin et al., 2017), and its increased tumor burden in AhR-lacking mice involves the expansion of undifferentiated hepatic pluripotent cells (Moreno-Marin et al., 2017) that unusually remain in the adult liver even after age-dependent maturation (Moreno-Marin et al., 2018). Notably, aging induced a significant increase in spontaneous (e.g., not induced) liver tumors in AhR deficient mice that was accompanied by larger numbers of liver progenitor cells eventually contributing to tumorigenesis as they do to improve hepatic regeneration (Mitchell et al., 2006; Moreno-Marin et al., 2017). Since AhR expression and activation largely decrease with age in wild type mice, we suggest that: first, such partial deficiency in receptor levels may facilitate or account for the existence of a basal tumor load and a reduction in undifferentiated cells in those mice and, second, the remaining AhR expression could be still enough to restrain liver tumorigenesis decreasing its incidence as compared to fully depleted *AhR-/-* animals.

Cell senescence has been linked to physiological aging and to age-related diseases (Campisi and Robert, 2014; Chiche et al., 2020). Remarkably, senescence not only increases during *in vivo* reprogramming of tumors by OKSM pluripotency factors (Abad et al., 2013) but it also induces reprogramming *in vivo* through p16^Ink4a^ (Mosteiro et al., 2016; Mosteiro et al., 2018). Thus, evidences support a functional interaction between senescence and stemness that impacts tumor development and tissue regeneration (Campisi and d’Adda di Fagagna, 2007; Chiche et al., 2017; Hernandez-Segura et al., 2018). In agreement with that hypothesis, aging in AhR deficient mice results in higher levels of liver senescence with up-regulation of inducers p16^Ink4a^, p21^Cip1^, IL1 and TNFα. Moreover, AhR depletion also increased hepatic levels of SASP stimulating factors MCP-2 and MMP-12 and reduced those of SASP inhibitors FGF and HGF and VEGF. Since the secretory phenotype from senescent cells contributes to tumor development (Schosserer et al., 2017; Zeng et al., 2018) producing proinflammatory cytokines, growth factors, and extracellular matrix remodelers that facilitate the expansion of transformed cells, we suggest that the profile of SASP markers in *AhR-/-* liver could contribute to their enhanced hepatocarcinogenesis. Pluripotency and reprogramming inducers OCT4 and NANOG, both repressed in an AhR-dependent manner during liver tumorigenesis and regeneration (Moreno-Marin et al., 2017), were over-expressed in AhR-null livers in parallel to their enhanced senescence and tumor burden. We therefore propose that AhR is at the crossroad of signalling pathways that integrate stemness, senescence and tumor development during liver aging. ChIP data indicate that AhR transcriptionally regulates the expression of p16^Ink4a^, p21^Cip1^ and TNFα, thus supporting a plausible causal role of this receptor in senescence.

AhR expression modulates senescence in different cell types and organs including mesenchymal embryonic fibroblasts (Alexander et al., 1998). Using passaging as a model for *in vitro* aging, we have found that MEFs from *AhR-/-* mice are more susceptible to develop senescence as determined by their higher levels of SA-β-Gal, *p16^Ink4a^* and *p21^Cip1^*. These data again suggest that AhR inhibits senescence and, in fact, its increased expression in *AhR+/+* MEFs with passaging may reveal a cellular response to control such process. Indeed, the Bcl-2 family inhibitor Navitoclax (Zhu et al., 2016) exerts a senolytic effect eliminating senescent cells more efficiently in absence of AhR, with downregulation of SA-β-Gal, *p16^Ink4a^* and *p21^Cip1^*. AhR levels were reduced in *AhR+/+* MEFs, in agreement with the idea that reduced senescence correlates with receptor inhibition. Consistently, CDK4/6 inhibitor Palbociclib/PD0332991 (Yoshida et al., 2016) produced an opposed response with respect to Navitoclax expanding senescence and, in parallel, increasing AhR levels early during the *in vitro* aging process. Interestingly, the effects of AhR on embryonic senescence were also seen in adult aged fibroblasts from *AhR-/-* mouse tail, in which cell cycle blockade and higher levels of SA-β-Gal, *p16^Ink4a^* and *p21^Cip1^* correlated with senescence intensity. Further, aged *AhR-/-* adult fibroblasts had decreased mitochondrial activity that could result from their enhanced senescence and a more pronounced aged phenotype as previously suggested (Moro, 2019; Picca et al., 2019). Finally, elimination of senescent cells *in vivo* by activation of p16^Ink4a^ in *p16^Ink4a^-3MR* transgenic mice further support our conclusion that lower senescence rates are associated to elevated AhR expression levels.

In summary, we provide here experimental evidence that the aryl hydrocarbon receptor is a molecular intermediate in the signalling pathways that govern the interaction between senescence and reprogramming. AhR would serve as a limiting factor in the control of both senescence and pluripotency eventually restricting tumor development with aging. Physiological modulation of AhR activity may represent a potential approach to circumvent progression of liver tumors in humans.

## METHODS

### Mice and treatments

*AhR+/+* and *AhR-/-* mice (C57BL/6N x 129 sV) were produced by homologous recombination in embryonic stem cells as described previously (Fernandez-Salguero et al., 1995) and used at 4-6 weeks (young group) or 15-22 months (aged group). For AhR induction, mice were injected with 4 µg/kg 6-formylindolo[3,2-b]carbazole (FICZ) diluted in PBS. Untreated control mice were injected with the same volume of DMSO. At the end of treatments, mice were sacrificed, and their livers extracted. *p16^Ink4a^-3MR* transgenic mice were generated as reported (Demaria et al., 2014). Mice at 4-7 months of age were treated i.p. with a single dose of 10 mg/kg doxycycline and seven days later with 25 mg/kg ganciclovir once per day for 5 consecutive days to clear senescent cells. Control mice received the same volume of vehicle (PBS). Experiments using mice were performed in accordance with the National and European legislation (Spanish Royal Decree RD53/2013 and EU Directive 86/609/CEE as modified by 2003/65/CE, respectively) for the protection of animals used for research. Experiments using mice were approved by the Bioethics Committee for Animal Experimentation of the University of Extremadura (Registry 109/2014) and by the Junta de Extremadura (EXP-20160506-1). Mice had free access to water and rodent chow.

### Mouse embryonic fibroblasts (MEFs)

*AhR+/+* and *AhR-/-* MEF cells were isolated from 14.5 d.p.c. mouse embryos as previously reported (Santiago-Josefat et al., 2001) and chemically induced senescence was performed as indicated (Guan et al., 2017). Briefly, MEFs cryopreserved for two days after isolation were thawed and grown in culture for 48 h in Dulbecco’s modified Eagle’s medium (DMEM) supplemented with 10% FBS, 2 mM L-glutamine, 100 U/ml penicillin and 100 mg/ml streptomycin at 37°C and 5% CO_2_ atmosphere. Cells were then trypsinized and plated at a density of 4 x10^5^ cells per 10-cm culture dish. For senescence induction, cultures were treated with 4 μM of the CDK4/6 inhibitor Palbociclib/PD-0332991 (Merck, PZ-0199) for 8 days, refreshing media and drug on day 4. Treated cells were trypsinized, washed with PBS and plated in complete DMEM medium for at least 24 h before starting the experiments. For depletion of senescent cells, MEFs were treated for 48 h with 10 μM of the Bcl-2 inhibitor Navitoclax (MedChemExpress, HY-10087) as described (Zhu et al., 2016). In both treatments, equivalent volumes of solvent were used as vehicle controls.

### Adult primary fibroblasts

Adult primary fibroblasts were obtained from tail tips of *AhR+/+* and *AhR-/-* mice. Mice were sacrificed and biopsies of the last 3 cm of tail were excised, cleaned in PBS and disinfected in 70% ethanol. After drying, tissues were sectioned in small pieces and placed in 0.25% trypsin solution (Invitrogen) in PBS and disaggregated by shaking during 90 min at 200 rpm and 37°C. The resulting digestion was filtered through a 40 μm mesh washing the remains of the digestion with RPMI medium. The filtrate was centrifuged at 200 g for 5 min and the resulting cell pellet was homogenized in RPMI medium containing 10% FBS, 50 μM 2-mercaptoethanol, non-essential amino acids (Gibco) and 1% penicillin/streptomycin at 37°C in a 5% CO_2_ atmosphere. At that time, fibroblast cultures were considered to be at passage 0. After 24 h, cultures were washed with PBS and fresh medium added. Thereafter, the medium was changed every other day.

### Gene expression analysis

Total RNA was isolated from *AhR+/+* and *AhR-/-* mouse livers using the Trizol reagent (Life Technologies). Following centrifugation, supernatants were precipitated with isopropanol, centrifuged again and pellets dissolved in DEPC-treated water. High Pure RNA Isolation Kit (Roche) was used to further purified RNA solutions from tissues and from *AhR+/+* and *AhR−/−* MEFs. To analyze mRNA expression by RT-qPCR, reverse transcription was performed using random priming and the iScript Reverse Transcription Super Mix (Bio-Rad). Real-time PCR was used to quantify the mRNA expression of *AhR*, *p16^Ink4a^, p21^Cip1^, IL1, TNFα, Tpr53, Mmp3, Cyp1a1* and *Gapdh* using SYBR® Select Master Mix (Life Technologies) in a Step One Thermal Cycler (Applied Biosystems) essentially as indicated (Rico-Leo et al., 2016). *Gadph* was used to normalize gene expression (ΔCt) and 2^−ΔΔCt^ to calculate changes in RNA levels with respect to control conditions. Primer sequences used are indicated in Supplementary Table 1.

### Immunoblotting

SDS–PAGE and immunoblotting were performed using total protein extracts obtained from *AhR+/+* and *AhR*-/- mouse livers as previously described (Rico-Leo et al., 2016). Briefly, tissues were extracted and homogenized in ice-cold lysis buffer; following centrifugation, protein concentration was determined in the supernatants using the Coomassie Plus protein assay reagent (Pierce) and bovine serum albumin (BSA) as standard. Aliquots of 20-30 µg total protein were electrophoresed in 8% SDS-PAGE gels which were transferred to nitrocellulose membranes. After blocking in a TBS-T (50 mM Tris-HC1 pH 7.5, 10 mM NaCl, 0.5% Tween 20) containing 5% non-fat milk, blots were sequentially incubated with the primary antibodies anti-AhR (Enzo 1:1000), anti-p53 (Cell signaling 1:1000) and anti-β-Actin (Sigma 1:1000) and secondary antibodies, washed in TBS-T and revealed using the Super-signal luminol substrate (Pierce). Blots were scanned and protein expression quantified in a ChemiDoc XRS+ equipment (Bio-Rad).

### Senescence-associated β-galactosidase activity

Senescence-associated β-galactosidase activity (SA-β-gal) was analyzed in primary fibroblasts, isolated liver cells and liver tissue following published protocols (Debacq-Chainiaux et al., 2009; Itahana et al., 2007). For primary adult fibroblasts and MEFs, the β-galactosidase fluorescent substrate C12FDG was added to the cultures at 60 µM concentration for 20 min and 37°C with gently shaking. C12FDG positive cells were quantified by flow cytometry using a Cytoflex S cytometer (Beckman Coulter) when analyzed in suspension, or with an Olympus FV1000 confocal microscope and the FV10 software (Olympus) when analyzed in attached cultures. In some experiments, MEFs were also stained with X-Gal as substrate to visualize the level of senescence by bright-field microscopy in a Olympus BX51 microscope. Isolated liver cells were obtained by mechanical disaggregation of liver tissue in a gentleMACS Octo Dissociator (Miltenyi). The resulting cell suspension was filtered through a 30 µm mesh and centrifuged for 5 min at 1800 g. Cell pellets were resuspended in PBS and incubated with the C12FDG substrate as indicated above. Cells were then resuspended in 0.1% BSA containing 10 µg/µl Hoechst 33258 and analyzed for senescence in a Cytoflex S cytometer (Beckman Coulter). For liver tissue, once fixed in buffered formaldehyde and included in paraffin, 3-5 μm liver sections were deparaffinized, rehydrated and stained for SA-β-Gal activity using X-Gal as substrate as described (de Mera-Rodriguez et al., 2019). Sections were observed and photographed using an Olympus BX51 microscope.

### Hematoxylin/eosin staining of liver sections

Mouse livers were processed for H&E staining as described (Moreno-Marín et al., 2018). Tissues were fixed overnight at room temperature in buffered formalin and included in paraffin. Sections of 3 µm were deparaffinated, rehydrated and incubated for 3 min with hematoxylin; after washing with tap water, they were stained with eosin for 1 min. Sections were then dehydrated, mounted and observed in a NIKON TE2000U microscope using 4x (0.10 numeric aperture), 10x (0.25 numeric aperture) an 20x (0.40 numeric aperture) objectives.

### Immunofluorescence

Liver sections were manually deparaffinated and rehydrated in PBS. Antigen unmasking was performed in citrate buffer at pH 6. After washing in PBS containing 0.05% Triton X-100 (PBS-T), non-specific epitopes were blocked by incubation for 1 h at room temperature in PBS-T containing 0.2% gelatin and 3% BSA (PBS-T-G-B). Sections were incubated overnight at 4°C with the following primary antibodies diluted in PBS-T-G-B: anti-p16^Ink4a^ (Santa Cruz Biotechnology 1:100), anti-p21^Cip1^ (Abcam, 1:100), anti-α-SMA (Sigma 1:200) anti-OCT4 (Santa Cruz Biotechnology 1:200) and anti-NANOG (Novus Biologicals 1:100). Following washing in PBS-T, sections were incubated for 1 h at room temperature with Alexa-488, Alexa-550 or Alexa-633 labeled secondary antibodies diluted in PBS-T-G-B. After additional washing, sections were dehydrated and mounted in PBS:glicerol (1:1). Primary fibroblasts were grown in glass coverslips, washed with PBS and fixed for 10-15 min at room temperature with 3.5% paraformaldehyde in PBS. After washing, cells were permeabilized by incubation in PBS containing 0.1% Triton X-100 and 0.2% BSA, washed again in PBS and blocked for 30 min in 2% BSA in PBS. Specific primary antibodies anti-p16^Ink4a^ (Novus Biologicals 1:200), anti-p21^Cip1^ (Santa Cruz Biotechnology 1:100) and anti-Cyclin E (Santa Cruz Biotechnology 1:75) were added in blocking solution overnight at 4°C. Following three washes with PBS, secondary antibodies labeled with Alexa-488 or Alexa-633 were added for 1 h. Liver and fibroblast samples were visualized using an Olympus FV1000 confocal microscope (Olympus). Fluorescence analysis was done using the FV10 software (Olympus). DAPI was used to stain cell nuclei.

### Mitochondrial membrane potential

Tail tip fibroblasts were seeded in 96-well plates and the mitochondrial membrane potential (MMP) was determined using the mitochondrial membrane potential kit (Sigma) following the recommendations made by the manufacturer. The red/green fluorescence intensity ratio was used to determine MMP values. To determine mitochondrial activity, fibroblasts were stained with 4 nM tetramethylrodamine for 30 min in complete RPMI medium without phenol red at 37°C and 5% CO_2_. Samples were analyzed using an Olympus FV1000 confocal microscope and the FV10 software (Olympus). Additional determination of MMP was done using the JC-10 dye. Fibroblasts were processed as indicated above and incubated with JC-10 for 30 min at 37°C and 5% CO_2._ Samples were analyzed using a Cytoflex S flow cytometer (Beckman Coulter).

### Chromatin Immunoprecipitation (ChIP)

Samples of liver tissue (60 mg) were finally minced, and DNA-protein interactions stabilized by incubation in 1% formaldehyde for 15 min at room temperature with agitation. Incubation was stopped by adding 0.125 glycine for 5 min. After washing in ice-cold PBS, samples were homogenized, and cell suspensions incubated for 10 min at 4°C in SDS lysis buffer containing protease inhibitors (Complete protease inhibitor cocktail, Roche). The remaining procedures were performed as described in previous reports (Carvajal-Gonzalez et al., 2009; Roman et al., 2008). Positive controls were done using input DNAs and negative controls were performed in absence of antibodies. Real-time PCR (qPCR) was caried out using IQ-SYBR Green in a Step One Thermal Cycler (Applied Biosystems). Specific primers used are indicated in Supplementary Table 2. Data are presented as percentage of DNA input in the antibody-containing immunoprecipitants minus the percentage of DNA input in the corresponding negative controls.

### Senescence associated secretory phenotype (SASP)

Liver tissue was homogenized in commercial lysis buffer (Bio-Rad) supplemented with protease inhibitors (Complete protease inhibitor cocktail, Roche) using a ratio of 1 mg tissue per 4 μl buffer. Samples were centrifuged at 500 g for 15 min and protein concentration determined in the supernatants. They were diluted with Bioplex sample diluent to a final concentration of 300 µg/ml. Blood plasma was obtained from whole blood taken from the left ventricle of the heart by cardiac puncture. Aliquots of 400-500 μl of blood were aspirated into a 2 ml syringe containing anticoagulant and immediately placed in EDTA containing tubes. Blood cells were separated by centrifugation at 1500 g for 15 min and the resulting plasma diluted 1:4 in Bioplex sample diluent. Cytokine content was analyzed using the Bio-Plex Multiplex immunoassay kit following the manufacturer’s instructions.

### Hexokinase activity assay

To determine glucose uptake by the liver, the Picoprobe Hexokinase activity kit (Biovision) was used following the manufacturer’s instructions. Samples of 30 mg of liver tissue were homogenized in 300 µl of cold HK assay buffer. After centrifugation for 5 min at 10.000 g and 4°C, supernatants were collected and used following the instructions provided by the manufacturer.

### Quantification of liver progenitor cells

Livers were removed from mice and processed using the gentleMACS Octo Dissociator kit (Miltenyi) to obtain homogeneous cell suspensions. Cell pellets were resuspended in 0.1% BSA in PBS and specific antibodies for undifferentiated liver cell markers CD133-PE (0.01 µg/µl) and CD326-EpCAM (0.3 µg/sample). A negative control without antibodies was used in each experimental condition. Cell suspensions were stirred for 20 min at 37°C and analyzed using a Cytoflex S Cytometer (Beckman Coulter).

### Statistical analyses

Quantitative data are shown as mean ± SD. Comparisons between experimental conditions was done using GraphPad Prism 6.0 software (GraphPad). The student’s t test was used to analyze differences between two experimental groups and ANOVA for the analyses of three or more groups. The Mann-Whitney non-parametric statistical method was used to compare rank variations between independent groups *(*P<0.05; **P<0.01, ***P<0.001)*, *\*\*\*\*P<0.0001)*.

## ACKNOWLEDGMENTS

We are very grateful to Dr. Javier de Francisco Morcillo (Departamento de Anatomía, Biología Celular y Zoología, Universidad de Extremadura) for his assistance in the analysis of SA-β-Gal expression. The authors acknowledge the support of the Servicio de Técnicas Aplicadas a las Biociencias (STAB-SAIUEX) de la Universidad de Extremadura.

## SUPPLEMENTARY MATERIAL

**Supplementary Figure 1.**
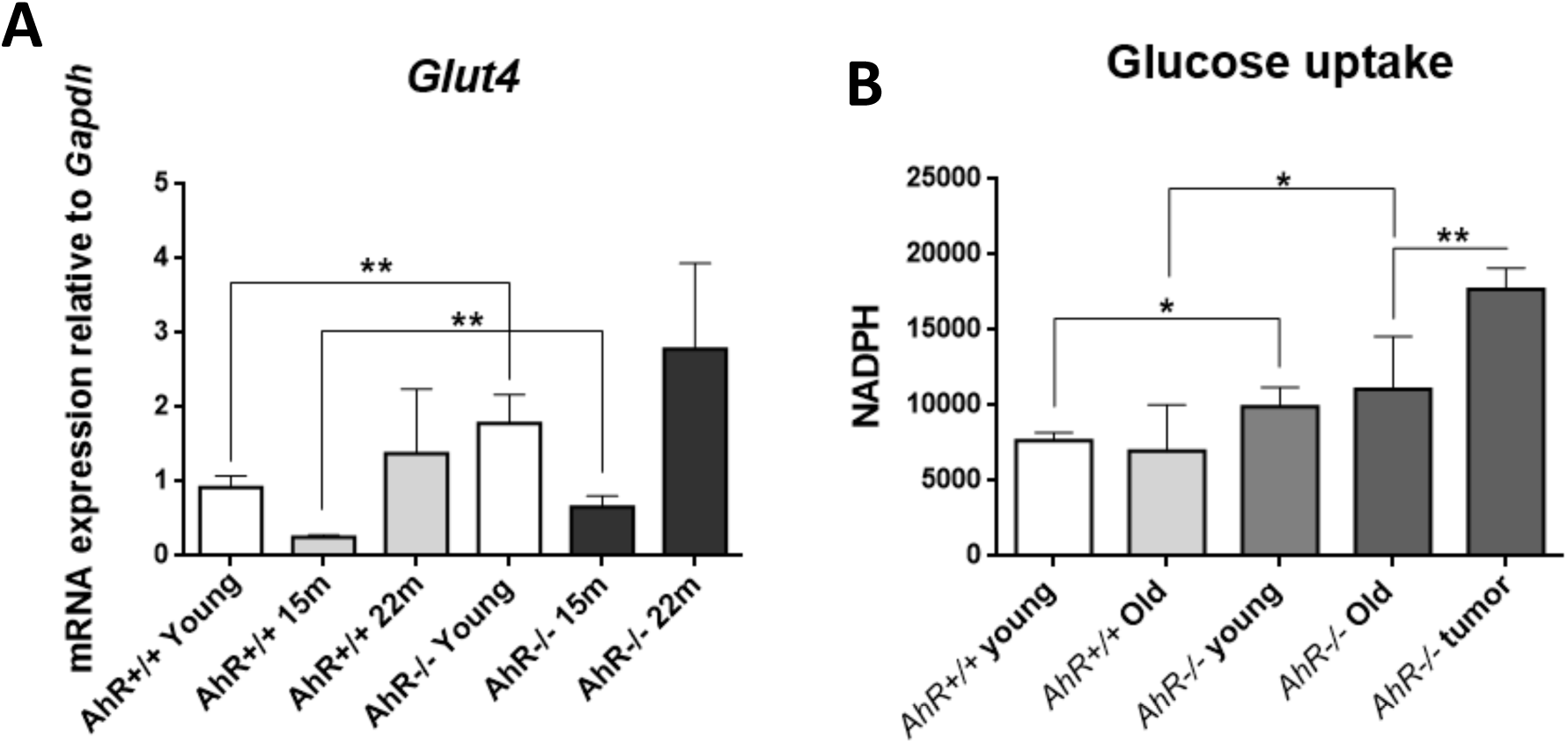
**(A)** mRNA expression of the glucose transporter *Glut4* was determined by RT-qPCR in *AhR+/+* and *AhR-/-* livers at the indicated ages using the oligonucleotides indicated in Supplementary Table 1. *Gapdh* was used to normalize target gene expression (ΔCt) and 2^−ΔΔCt^ to calculate changes in mRNA levels with respect to wild type or untreated conditions. **(B)** Glucose uptake was measured in young and aged *AhR+/+* and *AhR-/-* livers by the Picoprobe Hexokinase activity assay kit (*Biovision*). Data are shown as mean + SD. (\**P<0.05*; \*\**P<0.01*).

**Supplementary Table 1:** Oligonucleotide sequences used for mRNA expression analysis

**Supplementary Table 2:** Oligonucleotide sequences used for Chromatin immunoprecipitation (ChIP)

## REFERENCES

1. Abad, M., Mosteiro, L., Pantoja, C., Canamero, M., Rayon, T., Ors, I., Grana, O., Megias, D., Dominguez, O., Martinez, D., et al. (2013). Reprogramming in vivo produces teratomas and iPS cells with totipotency features. Nature 502, 340–345.

2. Alexander, D., Ganem, L., Fernandez-Salguero, P., Gonzalez, F., and Jefcoate, C. (1998). Aryl-hydrocarbon receptor is an inhibitory regulator of lipid synthesis and of commitment to adipogenesis. J Cell Sci 111, 3311–3322

3. Alvarez, D., Cardenes, N., Sellares, J., Bueno, M., Corey, C., Hanumanthu, V.S., Peng, Y., D’Cunha, H., Sembrat, J., Nouraie, M., et al. (2017). IPF lung fibroblasts have a senescent phenotype. Am J Physiol Lung Cell Mol Physiol 313, L1164–L1173.

4. Bravo-Ferrer, I., Cuartero, M.I., Medina, V., Ahedo-Quero, D., Pena-Martinez, C., Perez-Ruiz, A., Fernandez-Valle, M.E., Hernandez-Sanchez, C., Fernandez-Salguero, P.M., Lizasoain, I., et al. (2019). Lack of the aryl hydrocarbon receptor accelerates aging in mice. FASEB J 33, 12644–12654.

5. Brinkmann, V., Ale-Agha, N., Haendeler, J., and Ventura, N. (2019). The Aryl Hydrocarbon Receptor (AhR) in the Aging Process: Another Puzzling Role for This Highly Conserved Transcription Factor. Front Physiol 10, 1561.

6. Campisi, J. (2013). Aging, cellular senescence, and cancer. Annu Rev Physiol 75, 685–705.

7. Campisi, J., and d’Adda di Fagagna, F. (2007). Cellular senescence: when bad things happen to good cells. Nat Rev Mol Cell Biol 8, 729–740.

8. Campisi, J., and di Fagagna, F.d.A. (2007). Cellular senescence: when bad things happen to good cells. Nature reviews Molecular cell biology 8, 729.

9. Campisi, J., and Robert, L. (2014). Cell senescence: role in aging and age-related diseases. Interdiscip Top Gerontol 39, 45–61.

10. Carvajal-Gonzalez, J.M., Mulero-Navarro, S., Roman, A.C., Sauzeau, V., Merino, J.M., Bustelo, X.R., and Fernandez-Salguero, P.M. (2009). The dioxin receptor regulates the constitutive expression of the vav3 proto-oncogene and modulates cell shape and adhesion. Mol Biol Cell 20, 1715–1727.

11. Chiche, A., Chen, C., and Li, H. (2020). The crosstalk between cellular reprogramming and senescence in aging and regeneration. Exp Gerontol 138, 111005.

12. Chiche, A., Le Roux, I., von Joest, M., Sakai, H., Aguin, S.B., Cazin, C., Salam, R., Fiette, L., Alegria, O., Flamant, P., et al. (2017). Injury-Induced Senescence Enables In Vivo Reprogramming in Skeletal Muscle. Cell stem cell 20, 407–414 e404.

13. Cho, Y.C., Zheng, W., and Jefcoate, C.R. (2004). Disruption of cell-cell contact maximally but transiently activates AhR-mediated transcription in 10T1/2 fibroblasts. Toxicol Appl Pharmacol 199, 220–238.

14. Coppe, J.P., Patil, C.K., Rodier, F., Sun, Y., Munoz, D.P., Goldstein, J., Nelson, P.S., Desprez, P.Y., and Campisi, J. (2008). Senescence-associated secretory phenotypes reveal cell-nonautonomous functions of oncogenic RAS and the p53 tumor suppressor. PLoS Biol 6, 2853–2868.

15. Coutu, D.L., and Galipeau, J. (2011). Roles of FGF signaling in stem cell self-renewal, senescence and aging. Aging (Albany NY) 3, 920–933.

16. de Mera-Rodriguez, J.A., Alvarez-Hernan, G., Ganan, Y., Martin-Partido, G., Rodriguez-Leon, J., and Francisco-Morcillo, J. (2019). Senescence-associated beta-galactosidase activity in the developing avian retina. Dev Dyn 248, 850–865.

17. Debacq-Chainiaux, F., Erusalimsky, J.D., Campisi, J., and Toussaint, O. (2009). Protocols to detect senescence-associated beta-galactosidase (SA-betagal) activity, a biomarker of senescent cells in culture and in vivo. Nat Protoc 4, 1798–1806.

18. Demaria, M., Ohtani, N., Youssef, S.A., Rodier, F., Toussaint, W., Mitchell, J.R., Laberge, R.M., Vijg, J., Van Steeg, H., Dolle, M.E., et al. (2014). An essential role for senescent cells in optimal wound healing through secretion of PDGF-AA. Dev Cell 31, 722–733.

19. Eckers, A., Jakob, S., Heiss, C., Haarmann-Stemmann, T., Goy, C., Brinkmann, V., Cortese-Krott, M.M., Sansone, R., Esser, C., Ale-Agha, N., et al. (2016). The aryl hydrocarbon receptor promotes aging phenotypes across species. Sci Rep 6, 19618.

20. Fan, Y., Boivin, G.P., Knudsen, E.S., Nebert, D.W., Xia, Y., and Puga, A. (2010). The aryl hydrocarbon receptor functions as a tumor suppressor of liver carcinogenesis. Cancer Res 70, 212–220.

21. Fernandez-Salguero, P., Pineau, T., Hilbert, D.M., McPhail, T., Lee, S.S., Kimura, S., Nebert, D.W., Rudikoff, S., Ward, J.M., and Gonzalez, F.J. (1995). Immune system impairment and hepatic fibrosis in mice lacking the dioxin-binding Ah receptor. Science 268, 722–726.

22. Gorgoulis, V., Adams, P.D., Alimonti, A., Bennett, D.C., Bischof, O., Bishop, C., Campisi, J., Collado, M., Evangelou, K., Ferbeyre, G., et al. (2019). Cellular Senescence: Defining a Path Forward. Cell 179, 813–827.

23. Guan, X., LaPak, K.M., Hennessey, R.C., Yu, C.Y., Shakya, R., Zhang, J., and Burd, C.E. (2017). Stromal Senescence By Prolonged CDK4/6 Inhibition Potentiates Tumor Growth. Mol Cancer Res 15, 237–249.

24. Guerrina, N., Traboulsi, H., Eidelman, D.H., and Baglole, C.J. (2018). The Aryl Hydrocarbon Receptor and the Maintenance of Lung Health. International journal of molecular sciences 19, 3882.

25. Hernandez-Segura, A., Nehme, J., and Demaria, M. (2018). Hallmarks of Cellular Senescence. Trends Cell Biol 28, 436–453.

26. Itahana, K., Campisi, J., and Dimri, G.P. (2007). Methods to detect biomarkers of cellular senescence: the senescence-associated beta-galactosidase assay. Methods Mol Biol 371, 21–31.

27. Ito, T., Tsukumo, S., Suzuki, N., Motohashi, H., Yamamoto, M., Fujii-Kuriyama, Y., Mimura, J., Lin, T.M., Peterson, R.E., Tohyama, C., et al. (2004). A constitutively active arylhydrocarbon receptor induces growth inhibition of jurkat T cells through changes in the expression of genes related to apoptosis and cell cycle arrest. J Biol Chem 279, 25204–25210.

28. Kaiser, H., Parker, E., and Hamrick, M.W. (2020). Kynurenine signaling through the aryl hydrocarbon receptor: Implications for aging and healthspan. Exp Gerontol 130, 110797.

29. Ko, C.I., and Puga, A. (2017). Does the Aryl Hydrocarbon Receptor Regulate Pluripotency? Curr Opin Toxicol 2, 1–7.

30. Ko, C.I., Wang, Q., Fan, Y., Xia, Y., and Puga, A. (2013). Pluripotency factors and Polycomb Group proteins repress aryl hydrocarbon receptor expression in murine embryonic stem cells. Stem Cell Res 12, 296–308.

31. Kruk, J., Aboul-Enein, B.H., Bernstein, J., and Gronostaj, M. (2019). Psychological Stress and Cellular Aging in Cancer: A Meta-Analysis. Oxid Med Cell Longev 2019, 1270397.

32. Kuo, K.K., Lee, K.T., Chen, K.K., Yang, Y.H., Lin, Y.C., Tsai, M.H., Wuputra, K., Lee, Y.L., Ku, C.C., Miyoshi, H., et al. (2016). Positive Feedback Loop of OCT4 and c-JUN Expedites Cancer Stemness in Liver Cancer. Stem Cells.

33. Mitchell, K.A., Lockhart, C.A., Huang, G., and Elferink, C.J. (2006). Sustained aryl hydrocarbon receptor activity attenuates liver regeneration. Mol Pharmacol 70, 163–170.

34. Morales-Hernandez, A., Gonzalez-Rico, F.J., Roman, A.C., Rico-Leo, E., Alvarez-Barrientos, A., Sanchez, L., Macia, A., Heras, S.R., Garcia-Perez, J.L., Merino, J.M., et al. (2016). Alu retrotransposons promote differentiation of human carcinoma cells through the aryl hydrocarbon receptor. Nucleic Acids Res 44, 4665–4683.

35. Morales-Hernandez, A., Nacarino-Palma, A., Moreno-Marin, N., Barrasa, E., Paniagua-Quinones, B., Catalina-Fernandez, I., Alvarez-Barrientos, A., Bustelo, X.R., Merino, J.M., and Fernandez-Salguero, P.M. (2017). Lung regeneration after toxic injury is improved in absence of dioxin receptor. Stem Cell Res 25, 61–71.

36. Moreno-Marin, N., Barrasa, E., Morales-Hernandez, A., Paniagua, B., Blanco-Fernandez, G., Merino, J.M., and Fernandez-Salguero, P.M. (2017). Dioxin Receptor Adjusts Liver Regeneration After Acute Toxic Injury and Protects Against Liver Carcinogenesis. Sci Rep 7, 10420.

37. Moreno-Marin, N., Merino, J.M., Alvarez-Barrientos, A., Patel, D.P., Takahashi, S., Gonzalez-Sancho, J.M., Gandolfo, P., Rios, R.M., Munoz, A., Gonzalez, F.J., et al. (2018). Aryl Hydrocarbon Receptor Promotes Liver Polyploidization and Inhibits PI3K, ERK, and Wnt/beta-Catenin Signaling. iScience 4, 44–63.

38. Moreno-Marín, N., Merino, J.M., Alvarez-Barrientos, A., Patel, D.P., Takahashi, S., González-Sancho, J.M., Gandolfo, P., Rios, R.M., Muñoz, A., and Gonzalez, F.J. (2018). Aryl hydrocarbon receptor promotes liver polyploidization and inhibits PI3K, ERK, and Wnt/β-catenin signaling. iScience 4, 44–63.

39. Moro, L. (2019). Mitochondrial Dysfunction in Aging and Cancer. J Clin Med 8.

40. Mosteiro, L., Pantoja, C., Alcazar, N., Marion, R.M., Chondronasiou, D., Rovira, M., Fernandez-Marcos, P.J., Munoz-Martin, M., Blanco-Aparicio, C., Pastor, J., et al. (2016). Tissue damage and senescence provide critical signals for cellular reprogramming in vivo. Science 354.

41. Mosteiro, L., Pantoja, C., de Martino, A., and Serrano, M. (2018). Senescence promotes in vivo reprogramming through p16(INK)(4a) and IL-6. Aging Cell 17.

42. Mulero-Navarro, S., and Fernandez-Salguero, P.M. (2016). New Trends in Aryl Hydrocarbon Receptor Biology. Front Cell Dev Biol 4, 45.

43. Mulero-Navarro, S., Pozo-Guisado, E., Perez-Mancera, P., Alvarez-Barrientos, A., Catalina-Fernandez, I., Hernandez-Nieto, E., Saenz-Santamaria, J., Martinez, N., Rojas, J., Sanchez-Garcia, I., et al. (2005). Immortalized mouse mammary fibroblasts lacking dioxin receptor have impaired tumorigenicity in a subcutaneous mouse xenograft model. J Biol Chem 280, 28731–28741.

44. Picca, A., Guerra, F., Calvani, R., Bucci, C., Lo Monaco, M.R., Bentivoglio, A.R., Coelho-Junior, H.J., Landi, F., Bernabei, R., and Marzetti, E. (2019). Mitochondrial Dysfunction and Aging: Insights from the Analysis of Extracellular Vesicles. Int J Mol Sci 20.

45. Puga, A., Ma, C., and Marlowe, J.L. (2009). The aryl hydrocarbon receptor cross-talks with multiple signal transduction pathways. Biochem Pharmacol 77, 713–722.

46. Puga, A., Tomlinson, C.R., and Xia, Y. (2005). Ah receptor signals cross-talk with multiple developmental pathways. Biochem Pharmacol 69, 199–207.

47. Regina, C., Panatta, E., Candi, E., Melino, G., Amelio, I., Balistreri, C.R., Annicchiarico-Petruzzelli, M., Di Daniele, N., and Ruvolo, G. (2016). Vascular ageing and endothelial cell senescence: Molecular mechanisms of physiology and diseases. Mech Ageing Dev 159, 14–21.

48. Rhinn, M., Ritschka, B., and Keyes, W.M. (2019). Cellular senescence in development, regeneration and disease. Development 146.

49. Rico-Leo, E.M., Moreno-Marin, N., Gonzalez-Rico, F.J., Barrasa, E., Ortega-Ferrusola, C., Martin-Munoz, P., Sanchez-Guardado, L.O., Llano, E., Alvarez-Barrientos, A., Infante-Campos, A., et al. (2016). piRNA-associated proteins and retrotransposons are differentially expressed in murine testis and ovary of aryl hydrocarbon receptor deficient mice. Open Biol 6.

50. Roman, A.C., Benitez, D.A., Carvajal-Gonzalez, J.M., and Fernandez-Salguero, P.M. (2008). Genome-wide B1 retrotransposon binds the transcription factors dioxin receptor and Slug and regulates gene expression in vivo. Proc Natl Acad Sci U S A 105, 1632–1637.

51. Roman, A.C., Carvajal-Gonzalez, J.M., Merino, J.M., Mulero-Navarro, S., and Fernandez-Salguero, P.M. (2018). The aryl hydrocarbon receptor in the crossroad of signalling networks with therapeutic value. Pharmacol Ther 185, 50–63.

52. Roman, A.C., Carvajal-Gonzalez, J.M., Rico-Leo, E.M., and Fernandez-Salguero, P.M. (2009). Dioxin receptor deficiency impairs angiogenesis by a mechanism involving VEGF-A depletion in the endothelium and transforming growth factor-beta overexpression in the stroma. J Biol Chem 284, 25135–25148.

53. Ryu, Y.S., Kang, K.A., Piao, M.J., Ahn, M.J., Yi, J.M., Bossis, G., Hyun, Y.M., Park, C.O., and Hyun, J.W. (2019). Particulate matter-induced senescence of skin keratinocytes involves oxidative stress-dependent epigenetic modifications. Exp Mol Med 51, 1–14.

54. Safe, S. (2001). Molecular biology of the Ah receptor and its role in carcinogenesis. Toxicol Lett 120, 1–7.

55. Safe, S., Lee, S.O., and Jin, U.H. (2013). Role of the aryl hydrocarbon receptor in carcinogenesis and potential as a drug target. Toxicol Sci 135, 1–16.

56. Santiago-Josefat, B., Mulero-Navarro, S., Dallas, S.L., and Fernandez-Salguero, P.M. (2004). Overexpression of latent transforming growth factor-{beta} binding protein 1 (LTBP-1) in dioxin receptor-null mouse embryo fibroblasts. J Cell Sci 117, 849–859.

57. Santiago-Josefat, B., Pozo-Guisado, E., Mulero-Navarro, S., and Fernandez-Salguero, P. (2001). Proteasome inhibition induces nuclear translocation and transcriptional activation of the dioxin receptor in mouse embryo primary fibroblasts in the absence of xenobiotics. Mol Cell Biol 21, 1700–1709.

58. Schmidt, J.V., Su, G.H.-T., Reddy, J.K., Simon, M.C., and Bradfield, C.A. (1996). Characterization of a murine Ahr null allele: Involvement of the Ah receptor in hepatic growth and development. Proc Natl Acad Sci U S A 93, 6731–6736.

59. Schosserer, M., Grillari, J., and Breitenbach, M. (2017). The Dual Role of Cellular Senescence in Developing Tumors and Their Response to Cancer Therapy. Front Oncol 7, 278.

60. Wang, B., Kohli, J., and Demaria, M. (2020). Senescent Cells in Cancer Therapy: Friends or Foes? Trends in cancer.

61. Yin, X., Zhang, B.H., Zheng, S.S., Gao, D.M., Qiu, S.J., Wu, W.Z., and Ren, Z.G. (2015). Coexpression of gene Oct4 and Nanog initiates stem cell characteristics in hepatocellular carcinoma and promotes epithelial-mesenchymal transition through activation of Stat3/Snail signaling. J Hematol Oncol 8, 23.

62. Yoshida, A., Lee, E.K., and Diehl, J.A. (2016). Induction of Therapeutic Senescence in Vemurafenib-Resistant Melanoma by Extended Inhibition of CDK4/6. Cancer Res 76, 2990–3002.

63. Zaher, H., Fernandez-Salguero, P.M., Letterio, J., Sheikh, M.S., Fornace, A.J., Jr., Roberts, A.B., and Gonzalez, F.J. (1998). The involvement of aryl hydrocarbon receptor in the activation of transforming growth factor-beta and apoptosis. Mol Pharmacol 54, 313–321.

64. Zeng, S., Shen, W.H., and Liu, L. (2018). Senescence and Cancer. Cancer Transl Med 4, 70–74.

65. Zhu, C., Ikemoto, T., Utsunomiya, T., Yamada, S., Morine, Y., Imura, S., Arakawa, Y., Takasu, C., Ishikawa, D., and Shimada, M. (2014). Senescence-related genes possibly responsible for poor liver regeneration after hepatectomy in elderly patients. J Gastroenterol Hepatol 29, 1102–1108.

66. Zhu, Y., Tchkonia, T., Fuhrmann-Stroissnigg, H., Dai, H.M., Ling, Y.Y., Stout, M.B., Pirtskhalava, T., Giorgadze, N., Johnson, K.O., Giles, C.B., et al. (2016). Identification of a novel senolytic agent, navitoclax, targeting the Bcl-2 family of anti-apoptotic factors. Aging Cell 15, 428–435.

67. Zorin, V., Zorina, A., Smetanina, N., Kopnin, P., Ozerov, I.V., Leonov, S., Isaev, A., Klokov, D., and Osipov, A.N. (2017). Diffuse colonies of human skin fibroblasts in relation to cellular senescence and proliferation. Aging (Albany NY) 9, 1404–1413.

