## Supplemental Table 1 and 2 for "Aryl Hydrocarbon Receptor Blocks Aging-Induced Senescence in the Liver and Fibroblast Cells"

Supplementary Table S1

*Oligonucleotide sequences used for mRNA expression analysis*

| Gene name | Sequence 5´- 3´ |
|---|---|
| ***AhR*** | Fw: AGCCGGTGCAGAAAACAGTAA<br>Rv: AGGCGGTCTAACTCTGTGTGT |
| ***Cyp1a1*** | Fw: ACAGACAGCCTCATTGAGCA<br>Rv:GGCTCCACGAGATAGCAGTT |
| ***Tnfα*** | Fw: CCACCACGCTCTTCTGTCTA<br>Rv: CTCCACTTGGTGGTTTGCTA |
| ***IL1*** | Fw: CGGGTGACAGTATCAGCAAC<br>Rv: GACAAACTTCTGCCTGACGA |
| ***Glut4*** | Fw: ATGACCAAGCCCTGAATCTG<br>Rv: CGTAGGATGTGGTGATGACG |
| ***p16*** | Fw: TACCCCGATTCAGGTGAT<br>Rv: TTGAGCAGAAGAGCTGCTACGT<br>Fw: GTCGCAGGTTCTTGGTCACT<br>Rv: CGAATCTGCACCGTAGTTGA |
| ***p21*** | Fw: GCCTTAGCCCTCACTCTGTG<br>Rv: AGCTGGCCTTAGAGGTGACA |
| ***p53*** | Fw: TGGAAGACTCCAGTGGGAA<br>Rv: TCTTCTGTACGGCGGTCTCT |
| ***Mmp3*** | Fw: AGTCAGGGTCACCCACAAAG<br>Rv: GCATTGGGTATCCATCCATC |
| ***Gapdh*** | Fw: TGAAGCAGGCATCTCAGGG<br>Rv: CGAAGGTGCAAGAGTGGGA |

Supplementary Table S2

*Oligonucleotide sequences used for Chromatin Immunoprecipitation (ChIP)*

| Gene name | Sequence 5´-- 3´ |
|---|---|
| ***p16*** | Fw: GGCACTCCCAGCAAGTAGAT<br>Rv: CACAAACGTGCCTCCTATACA |
| ***p21*** | Fw: TTTGTTGTCCTCGCCCTCAT<br>Rv: ACGCACGTACACAGACACA |
| ***Tnfα*** | Fw: CCAGACACTCACCTCATCCC<br>Rv: TGGAACTGGCAGAAGAGGC |
| ***Gapdh*** | Fw: TGAAGCAGGCATCTCAGGG<br>Rv: CGAAGGTGCAAGAGTGGGA |
